# Linear motif specificity in signaling through p38α and ERK2 mitogen-activated protein kinases

**DOI:** 10.1101/2022.08.23.505039

**Authors:** Jaylissa Torres Robles, Guangda Shi, Benjamin E. Turk

## Abstract

Mitogen-activated protein kinase (MAPK) cascades are essential for eukaryotic cells to integrate and respond to a wide array of stimuli. Maintaining specificity in signaling through MAPK networks is key to coupling specific inputs to appropriate cellular responses. One way that MAPKs achieve specificity is through transient interactions with docking sites: short linear motifs found in MAPK substrates, regulators, and scaffolds. Docking sites bind to a conserved groove located in the catalytic domain of all MAPKs including the ERK and p38 subfamilies, but how specificity is achieved remains unresolved. To understand the basis of docking selectivity for these two subfamilies, we screened a library of thousands of human proteome-derived sequences for docking to ERK2 and p38α. We discovered a large number of sequences that bound specifically to only one MAPK or promiscuously to both, and that selective and non-selective interactors conformed to distinct sequence motifs. In particular, selective binding to p38α correlated with higher net charge in the docking site, and this phenomenon was driven by enrichment for Lys residues. A pair of acidic residues unique to the docking groove of p38α mediated selectivity for Lys-rich basic motifs. Finally, we validated a set of full-length proteins harboring docking sites selected as hits in our screens to be authentic MAPK interactors and identified ChREBP and TACC1 as cellular MAPK substrates. This study identifies distinguishing features that help define MAPK signaling networks and explains how specific docking motifs promote signaling integrity.

## INTRODUCTION

Mitogen-activated protein kinases (MAPKs) are Ser/Thr kinases at the bottom of three-tiered kinase cascades that are essential for transmitting extracellular signals to effect changes in cell physiology and function^1^. Mammals have four canonical MAPK subfamilies, including the p38, c-Jun N-terminal kinase (JNK), extracellular signal-regulated kinase (ERK) 1/2, and ERK5 MAPKs. The various MAPK cascades respond to different stimuli and control distinct cellular processes^2-4^. For example, activation of the ERK pathway by growth factors promotes cell proliferation^2^, while the stress-activated p38 MAPKs are key to stress induced cell cycle arrest^5^ and pro-inflammatory signaling in post-mitotic cells^3^. Distinct biological roles played by each MAPK are in part mediated by connections with unique sets of interacting proteins including substrates, scaffolds and regulators. While numerous substrates and binding partners have been identified for each MAPK, their broad physiological and pathological functions suggest that key components of their signaling networks remain to be discovered. A more complete understanding of how MAPKs make selective connections with substrates and other interacting proteins would facilitate discovery of these components.

Numerous mechanisms can underlie signaling specificity by protein kinases, including subcellular compartmentalization, scaffolding, signal dynamics, and direct kinase-substrate interactions^6^. One mechanism common to Ser/Thr protein kinases involves the recognition of sequence motifs surrounding sites of phosphorylation by the catalytic cleft^7^. However, since all MAPKs target a common Ser/Thr-Pro consensus sequence, catalytic site specificity is insufficient to mediate selectivity for a particular member of the family. For MAPKs, docking interactions at regions outside of the catalytic cleft reportedly control specificity and impact their biological function^8,9^. In particular, MAPKs harbor a docking groove within the catalytic domain called the D-recruitment site (DRS). The DRS is an extended shallow surface comprising of several hydrophobic pockets and a negatively charged region termed the common docking (CD) region. The DRS binds to short linear motifs (SLiMs) known as D-sites (also called δ-domains or DEJL motifs), found in MAPK substrates and other interactors^10,11^. Like other SLiMs, D-sites are generally found in intrinsically disordered regions (IDRs) of proteins and bind to MAPKs transiently, with moderate to weak affinity, through a limited number of direct contacts. While these binding interactions are optimal for mediating substrate recruitment^12^, their transient nature makes them difficult to discover and as a result a small number of D-sites have been studied.

D-sites have been subcategorized into “forward” or “reverse” sequences as defined by their binding orientation in the DRS^13^. Forward D-sites, which are the most common, conform to a consensus sequence (β_2-3_-x_1-6_-ϕ-x-ϕ) consisting of a cluster of basic residues (β), connected to a hydrophobic motif (ϕ-x-ϕ) by a variable length linker (x) sequence^10,13-16^. Within the context of this general consensus, several sequence motifs with distinct linker amino acid composition and length, reportedly confer selective binding to particular MAPK subfamilies^17-21^. Structural studies of MAPKs in complex with D-site peptides (D-peptides) have revealed distinct binding modes associated with reported motifs, in some cases allowing rationalization of MAPK selectivity^13,15,22-29^. In particular, unique features of the JNK DRS facilitate substantial discrimination between cognate and non-cognate motifs^13,19,21^. While reverse D-sites peptides from the homologous kinases RSK1 and MK2 bind with a high degree of selectivity (>10-fold) to ERK and p38 MAPKs respectively, the extent to which forward D-sites distinguish between members of these two MAPK subfamilies is less clear^13,30^. In particular, specific features within D-sites conferring selective binding to their cognate MAPKs, and how their respective docking grooves encode specificity, are not known.

Here we define ERK2 and p38α D-site interactomes by screening a library of candidate docking sequences from the human proteome. These screens revealed distinct MAPK-selective D-site sequences and associated motifs. We show that motifs with Lys residues clustered in the D-site variable linker region conferred selectivity for p38α over ERK2. Furthermore, through exchange mutagenesis we identified a region of the DRS largely responsible for differential binding to the two MAPKs. In addition, our screen uncovered cryptic docking sequences in several established MAPK substrates and led us to discover a set of previously unknown ERK2 and p38α interactors and substrates. Overall, these studies reveal new connections in MAPK signaling networks and demonstrate how these connections occur selectively.

## RESULTS

### Establishment of a yeast two hybrid (Y2H) screening platform to identify MAPK D-sites

We established a yeast two-hybrid (Y2H) assay to probe the interaction between MAPKs and putative docking sequences, using a fragment of the MAPK substrate ELK1 (residues 306-427) including its essential D-site motif^31^ (Fig. 1A and Fig. S1). A Y2H reporter strain was co-transformed with a bait plasmid expressing the MAPK fused to the Gal4 DNA binding domain (Gal4^DBD^-ERK2 or Gal4^DBD^-p38α) and a prey plasmid expressing the ELK1 fragment fused to the Gal4 activation domain (Gal4^AD^-ELK1) (Fig. 1B) and assayed for growth under selective (His^-^) and non-selective (His^+^) conditions (Fig. 1B, C). We found that ELK1 could interact physically with ERK2, but not p38α (Fig. 1C, D). Importantly, mutation of the D-site (ELK1^ΔD^) disrupted binding, confirming the interaction to be D-site dependent. We next examined whether other cognate or non-cognate docking sequences could support this interaction. As anticipated based on reported peptide binding affinities^13,14,21^, replacement of the native ELK1 D-site with that of MAP2K2 (ELK1^MAP2K2-D^) allowed it to interact with both ERK2 and p38α. Conversely, substitution with a JNK-selective sequence (ELK1^NFAT4-D^) abolished binding to either kinase (Fig. 1C, D). Consistent with prior reports that a MAP2K2 D-site peptide binds more tightly to ERK2 than does that of ELK1, ELK1^MAP2K2-D^ imparted faster yeast growth than ELK1^WT^ when co-expressed with Gal4^DBD^-ERK2^14^. Taken together, this analysis suggests that Y2H is a suitable strategy to discover D-site sequences that bind competitively to MAPKs.

**Figure 1.**
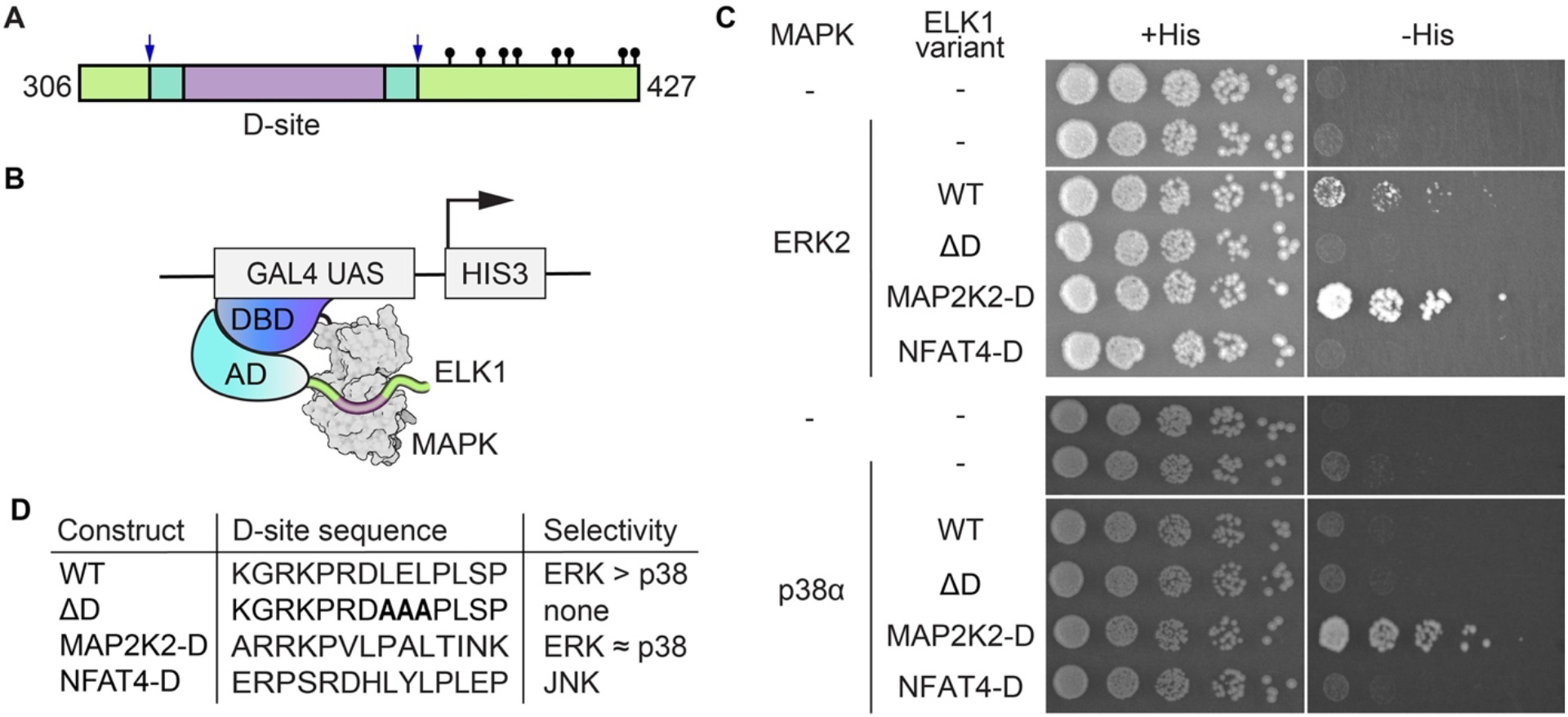
Y2H system to detect MAPK binding to docking sites. (A) Diagram of ELK1 transcriptional activation domain construct showing the D-site (purple) with flanking linker sequence (cyan) and MAPK Ser/Thr-Pro phosphorylation sites (lollipops). (B) Docking dependent yeast two-hybrid scheme. (C) Assays of yeast growth under selective (-His + 50 μM 3-AT) and non-selective conditions (+His) with strains co-expressing the indicated ELK1 variants with either ERK2 or p38α. The spot series was from five-fold serial dilutions of exponentially growing yeast cultures. (D) D-site sequences used in ELK1 constructs with their reported MAPK binding selectivity^13^.

### Y2H screen to define ERK2 and p38α D-site interactomes

To identify additional sequence features that mediate selective docking to ERK2 or p38α, we prepared a custom library of ELK1 D-site exchange mutants in the Gal4^AD^-ELK1 prey vector background. The library incorporated a set of 11,756 SLiM sequences from the human proteome conforming to the general forward D-site sequence motif [R/K]-x_0-2_-[R/K]-x_3-5_-[ILV]-x-[FILMV], which we have previously used in a different screening platform^19^ (Dataset S1). Yeast bearing Gal4^DBD^-MAPK bait plasmids were transformed with the prey plasmid library, and transformants were pooled and grown under selective and non-selective conditions in liquid culture. Cultures were periodically sampled, and the D-site coding sequence was PCR-amplified from extracted plasmid DNA and analyzed by next-generation sequencing (Fig. 2A). As expected, we saw significant enrichment of specific D-site sequences by ERK2 and p38α only under selective conditions (Fig. 2B, C and Fig. S2A-B). We assigned an enrichment score (ES) for each sequence by fitting the change in relative abundance over the first ∼15 population doublings to an exponential function. ES values for the full set of sequences correlated significantly in pairwise comparisons of three replicate screens for both ERK2 (range of Pearson’s correlation coefficient [*R*] = 0.60 – 0.74) and p38α (*R* = 0.82 – 0.85) and (Fig. S2C-D). Sequences with average ES values ≥ 2 standard deviations from the mean of all sequences (Z-score ≥2) were defined as hits. ERK2 and p38α hits (Datasets S2 and S3, Tables S1 and S2) included most previously known interacting sequences^15,17,23,32-35^ (Fig. 2D, E and Fig. S2B), though several established D-sites with low affinity for p38α escaped detection.

**Figure 2.**
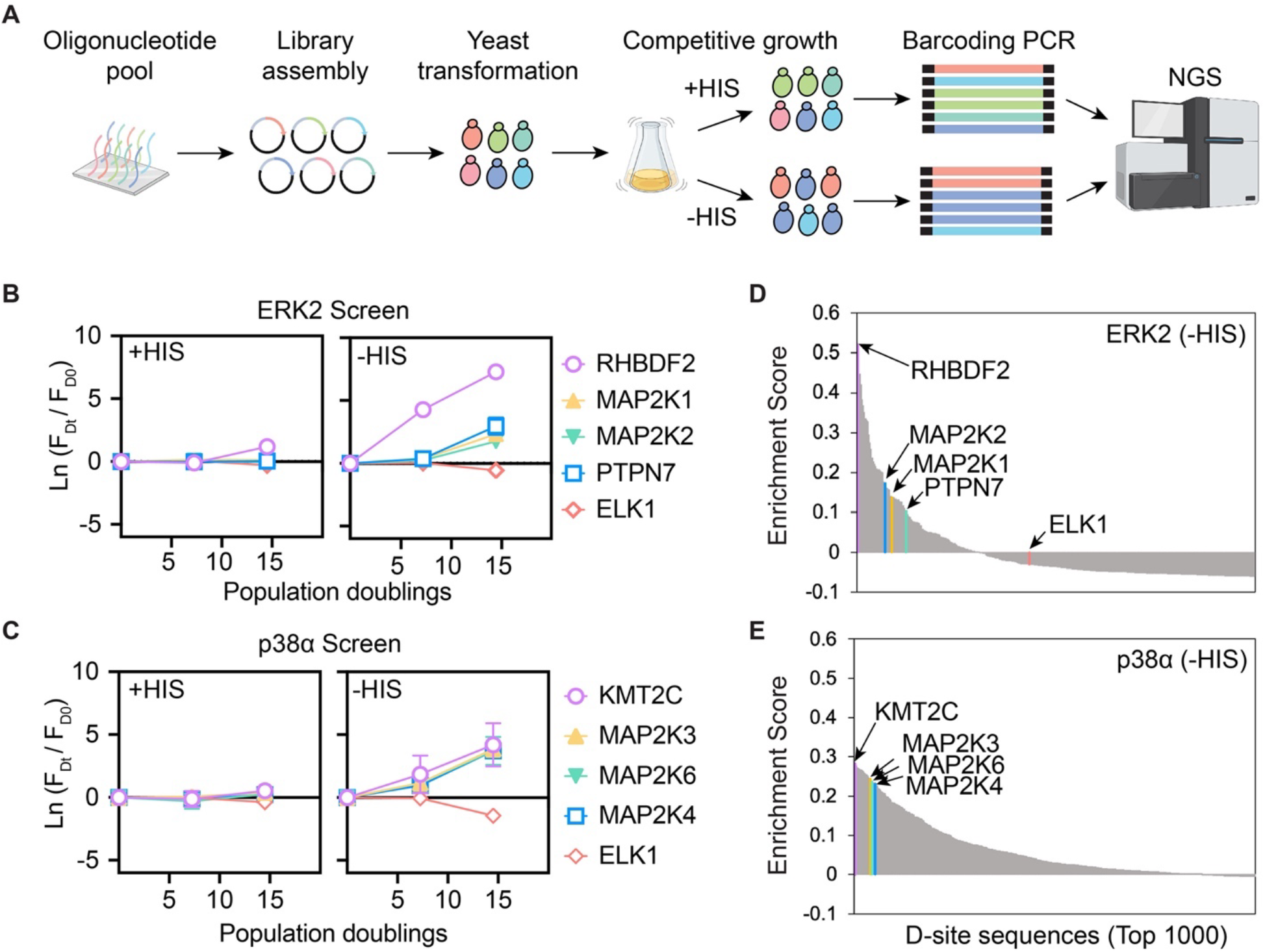
Pooled Y2H screening. (A) Library construction and Y2H screening workflow. (B, C) Natural logarithm (ln) of the change in frequency (F_Dt_/F_D0_) of selected library sequences in the ERK2 and p38α screens plotted as a function of yeast population doublings under non-selective (left panel) and selective (right panel) conditions. Shown are the most enriched sequences (RHBDF2 for the ERK2 screen and KMT2C for the p38α screen), several known interactors, and the parental ELK1 sequence. Error bars indicate SD from three separate screens. (D, E) Waterfall plots show average enrichment scores (ES) (in descending order, from three independent screens) for the 1000 most enriched sequences in the ERK2 and p38α screens under selective conditions. Specific interactors shown in (B) and (C) are highlighted.

Contrary to the generally good correlation between replicates for each MAPK, average ES values from ERK2 and p38α hits showed relatively poor correlation (*R* = 0.57), with many sequences enriched only in the presence of a single MAPK. Overall, screen hits included 98 sequences that were selected only by ERK2, 238 sequences selected only by p38α and 105 sequences that were hits in both screens (termed “common” hits from here on) (Fig. 3A, Tables S1 and S2). To validate the results from Y2H screens, we determined ERK2 and p38α binding affinities for a set of synthetic D-peptides corresponding to the top-ranking screening hits as well as several control sequences. Peptide binding was assayed by competitive inhibition of MAPK kinase activity using a D-site dependent fluorescent peptide substrate^36^. D-peptide IC_50_ values ranged from 50 nM to 100 µM for at least one MAPK (Figs. 3B, Fig. S3 and Table S3), consistent with previous studies indicating that D-sites bind to MAPKs with moderate affinity^13,20^. In general, we observe a trend for peptides to display higher affinity for p38α, including a number of common hits and a single hit sequence unique to the ERK2 screen (Fig. 3B, C and Table S3). These observations suggest that the ERK2 and p38α screens may have different affinity thresholds for significant enrichment in the context of the ELK1 sequence. It is also possible that p38α has a propensity to bind more tightly to D-site peptides than does ERK2, at least in the context of sequences present in our library. Overall, these results confirm the selection of true MAPK biding sequences by Y2H screens and delimits ERK2 and p38α D-site interactomes within the context of our library.

**Figure 3.**
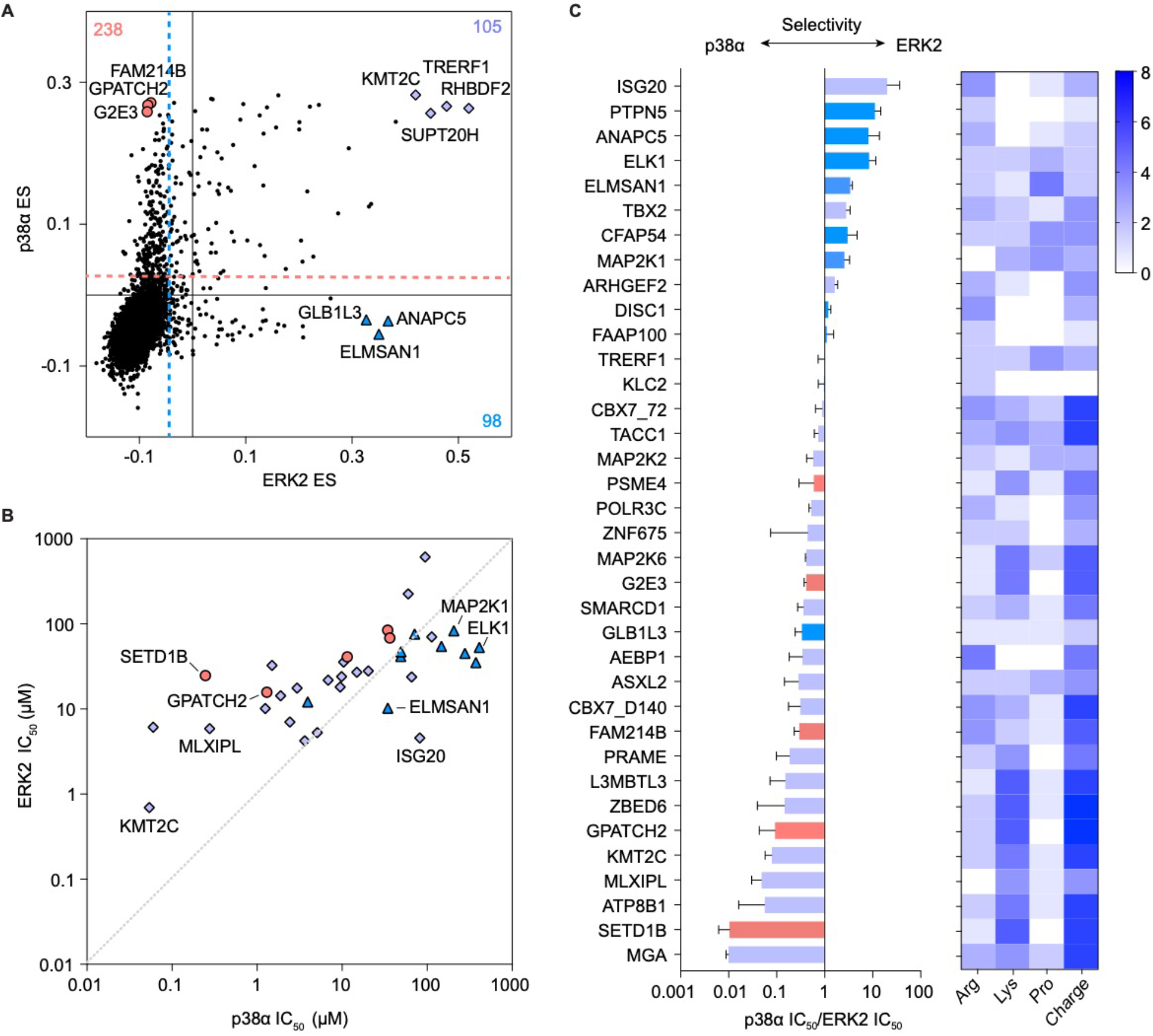
Comparison of ERK2 and p38α screens and hit validation. (A) The mean enrichment score (ES) is plotted for each sequence in the ERK2 (x-axis) and p38α (y-axis) Y2H screens. Dotted lines show the Z ≥2 hit threshold. The number of sequences selected uniquely by ERK2 (bottom-right quadrant), uniquely by p38α (upper-left quadrant), and by both MAPKs (upper-right quadrant) is indicated. (B) Plot shows mean IC_50_ values (from three independent experiments) for inhibition of ERK2 and p38α activity by synthetic D-peptides encoding hit D-site sequences. For selected peptides, the name of the corresponding gene is shown. ERK2-specific hits are shown as blue triangles, p38α-specific hits as red circles, and common hits are purple diamonds. (C) Bar graph shows selectivity ratios for D-peptides calculated from IC_50_ values in (B). D-peptide sequences and source data are provided in Table S3. Bars show mean ± SD from three independent experiments. The heat map shows the net peptide charge (at pH 7.0) and the abundance of Arg, Lys and Pro residues for each D-peptide.

### Identifying determinants of MAPK selectivity

We next analyzed our dataset to identify sequence features preferably recognized by each MAPK. Considered in aggregate, ERK2 and p38α both significantly selected sequences with Leu residues at positions 10 and 12, corresponding to the two residues that we required to be hydrophobic in the library design. In contrast, sequences selected by the two MAPKs differed substantially in residues over-represented at positions 7 – 9 and 13 (Fig. 4A, B). ERK2 was significantly selective for Pro at each of these positions (Fig. 4A), while p38α was significantly selective for basic residues and against acidic residues (Fig. 4B). These differences in amino acid composition correlated with a tendency for p38α hits to carry a higher net positive charge than those of ERK2 (Fig. 4C). Furthermore, in our validation set of synthetic D-peptides, highly charged and basic-rich D-peptides tend to have higher affinity for p38α over ERK2, while Proline-rich D-peptides tend to prefer ERK2 (Figs. 3C).

**Figure 4.**
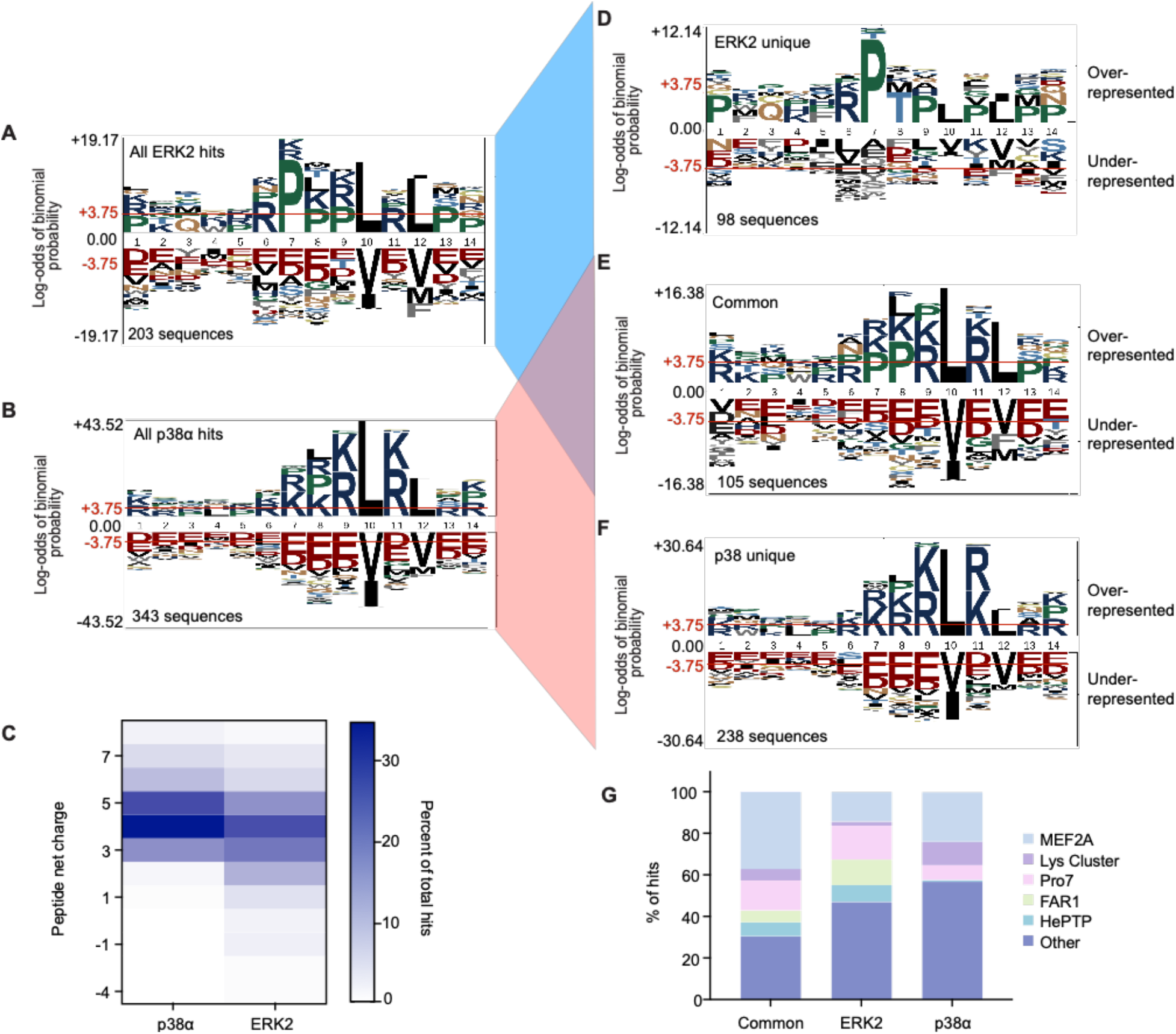
Sequence motifs selected by ERK2 and p38α. (A,B) Probability logos (pLogos)^77^ generated from multiple sequence alignments of all ERK2 or all p38α hits. (C) Heat map showing the percentage of total hits for each MAPK with a given calculated net charge at pH 7.0. (D-F) pLogos are shown corresponding to unique ERK2 hits, common hits, and unique p38α unique. (G) Distribution of motif classes for common and unique ERK2 and p38α hits.

Previous studies have defined multiple D-site motifs that bind non-selectively to both ERK and p38^13,20^. Thus, we mined aligned hit sequences from our screens to identify exclusive motif classes. When we subdivided hits into those either unique to a single MAPK or common to both, it became evident that sequences in each category conform to distinct motif signatures (Fig. 4D-G). For example, 30% of hits unique to ERK2 had a Pro residue at position 7 (P7), and more than half of these sequences were significantly enriched for an additional Pro exclusively at position 9 and were devoid of basic residues at position 11 (Fig. 4D and Fig. S4). The full P-x-P-L-x-ϕ motif, which corresponded to 12% of ERK2 unique hits, has been previously defined as the “Far1” motif^17,22^. ERK2 unique hits lacking Pro at position 7 (14%) generally conformed to the previously defined “MEF2A” motif class characterized by aliphatic residues at position 8 (Fig. 4G and Fig. S4)^17^. Notably, the MEF2A motif was also found in a substantial proportion (24%) of p38α unique hits (Fig. 4G and Fig. S5), and it was the dominant signature in hits common to both MAPKs (Fig. 4E, G). However, unlike its prior definition (P/ϕ-x-L-x-L), we found MEF2A class sequences to be significantly enriched for basic residues at positions 9 and 11 (Fig. 4E, Fig. S4 and Fig. S5). In contrast to ERK2 hits, unique p38α hits had basic residues clustered at positions 7 – 9 as a predominant feature (Fig. 4F). At these positions we observed significant over-representation of Lys but not Arg residues, defining a previously unknown motif (K_2-3_-L-x-ϕ) which we termed “Lys cluster” (Fig. S5C). Notably, of the nine sequences in our library having three Lys residues at positions (7 – 9), seven scored as p38α hits. Consistent with these observations, the effect of net positive charge on relative affinity of synthetic D-peptides for p38α over ERK2 correlated well with the number of Lys residues (Fig. 3C). Finally, a number of p38α (2%) and ERK2 (6%) hits included an L/I/V-x-x-R-R sequence motif near the N-terminus of the D-site, a motif previously noted in the D-site from the phosphatase PTPN7 (HePTP) with few other known examples (Fig. 4C)^17,23^. Overall, through these analyses we were able to identify hits containing previously known MAPK docking motifs and an unknown signature promoting selective binding to p38α over ERK2.

Among the defined motifs, the Far1 and Lys cluster motifs exhibited the largest differences in representation between ERK and p38 hits (Fig. 4G). To assess the extent to which these sequence motifs interact selectively with ERK2 and p38α, we generated two D-peptides (ERK2-D-pep and p38α-D-pep) incorporating residues specifically selected by the corresponding MAPKs in the context of the Far1 and Lys cluster motifs, respectively. We determined their relative affinity for binding to ERK2 and p38α by competitive kinase assay as described above (Fig. 5A, Table S3). Both peptides bound their corresponding MAPKs with relatively high affinity (IC_50_ values ∼2 μM), likely due to inclusion of favorable residues throughout the peptide sequence. Consistent with their anticipated selectivity, p38-D-pep bound p38α with 10-fold higher affinity than ERK2, while ERK-D-Pep showed a more modest 2.4-fold preference for ERK2. We also evaluated a pair of consensus peptides designed in the context of the common MEF2A motif (ERK2-MEF-pep and p38α-MEF-pep) that differ in estimated net charge (6 and 8 respectively, at neutral pH). We found that ERK2-MEF-pep bound ERK2 and p38α with similar affinities, while p38-MEF-pep favored p38α by 9-fold (Fig. 5A). These results with MEF2A motif derived peptides confirm that enrichment for basic residues and high net charge correlate with p38α selectivity. Altogether these observations confirm that motifs identified by multiple sequence alignment of Y2H screen hits confer preferential binding to p38α and to a lesser extent ERK2.

**Figure 5.**
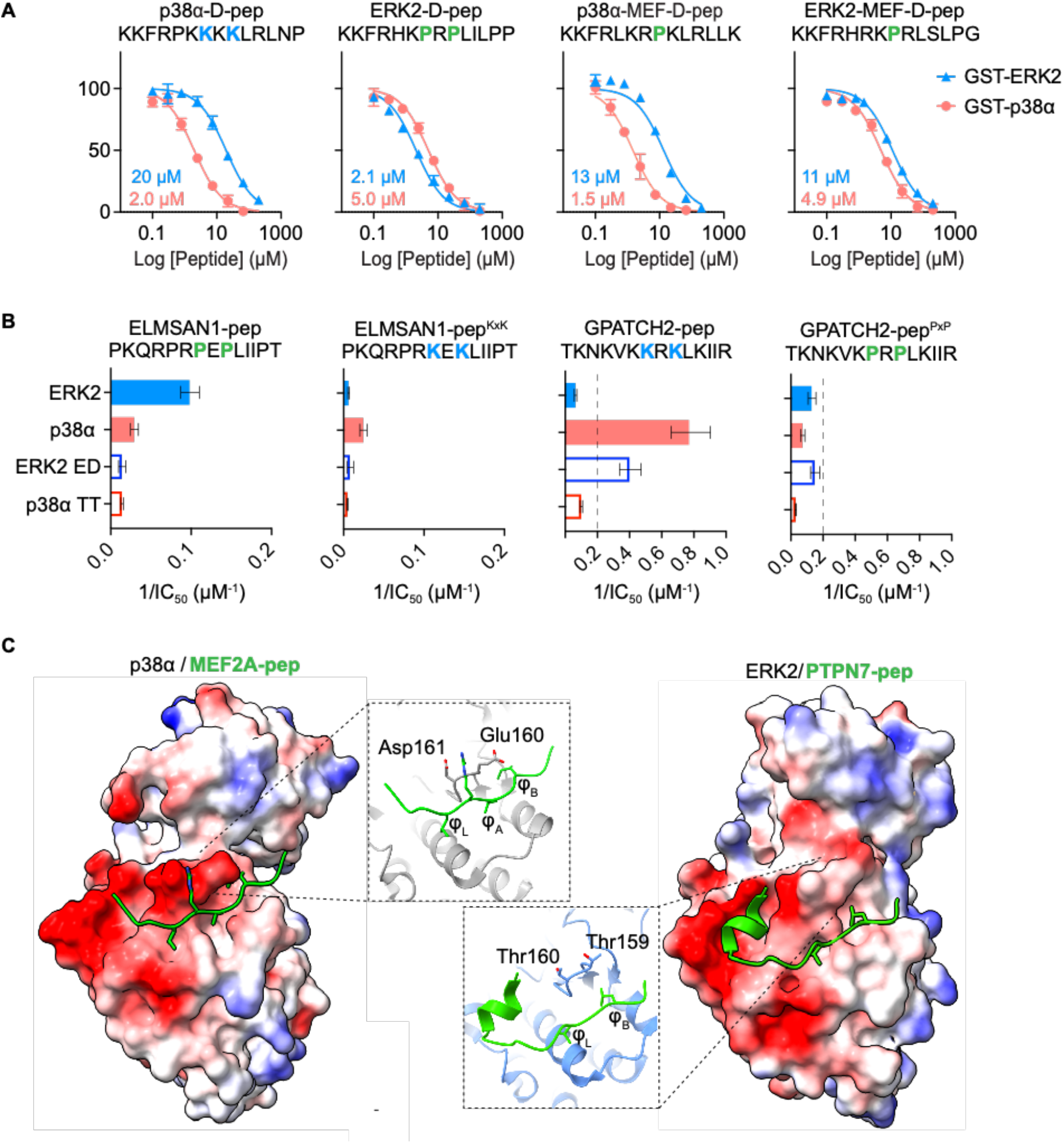
Validation of MAPK-selective D-site motifs. (A) Dose response curves for inhibition of ERK2 (blue, triangle) or p38α (red, circle) by the indicated synthetic D-peptides. Error bars represent standard deviations (SD) from three independent experiments. (B) Inverse IC_50_ values for inhibition of ERK2, p38α, ERK2^T159E/T160E^, or p38^E160T/D161T^ by indicated synthetic peptides. Error bars represent 95% confidence intervals from three experiments. (C) Structures of p38α (left, PDB: 1LEW^15^) and ERK2 (right, PDB: 2GPH^23^) in complex with D-peptides, showing electrostatic potentials calculated in ChimeraX^79^. Surface coloring represents columbic values. The insets show interactions of MEF2A and PTPN7 D-peptides (green) with p38α and ERK2, respectively. The ED site residues are labeled and shown in stick representation.

To systematically test the capacity of Lys- and Pro-rich motifs to specifically bind p38α and ERK2 respectively, we examined the impact of eliminating or exchanging these features in the context of p38 selective (GPATCH2-pep, with a native KxK motif) and ERK selective (ELMSAN1-pep, with a native PxP motif) D-peptides. As anticipated, substitution of the key residues with Ala in these peptides led to large reduction in binding affinity to both MAPKs (Fig. 5B, Fig. S6A, B and Table S3). Moreover, introduction of Pro residues into GPATCH2-pep (GPATCH2-pep^PxP^) caused a >10-fold reduction in affinity for p38α and 2-fold increased affinity for ERK2 (Fig. 5B, Table S3). The converse substitutions in ELMSAN1-pep (ELMSAN1-pep^KxK^) provided >100-fold reduced affinity towards ERK2 without significant change in affinity for p38α. These results confirm the trends seen with the hit-derived consensus D-peptides that though Pro-rich sequences bind with modestly lower IC_50_ values towards ERK, Pro residues are still permissive for binding to p38α. Finally, these results suggest that basic residues clustered in positions 7 - 9 promote specificity for p38α and disrupt ERK2 binding.

To rationalize these findings, we examined previously determined co-crystal structures of p38α and ERK2 in complex with D-peptides. Consistent with its enhanced selectivity for peptides with high net charge, the p38α DRS has an extended negative charge distribution across the entire DRS groove in comparison to ERK2 (Fig. 5C). Negative charge in the p38α groove is concentrated in two areas: the CD region (E81, D316 and E317 in loop L16) and the so-called “ED site” localized to the β7-β8 turn (E160 and D161). ERK2 also harbors a negatively-charged CD region (E81, D320 and E321), but has a pair of neutral residues (Thr159 and Thr160) in place of the ED site (Fig. 5C). The β7-β8 turn residues have been implicated in selective binding of reverse D-sites to p38 and ERK^15,25,34,35,37-39^. We hypothesized that the extended negative surface potential provided by ED site promotes p38α specificity for forward D-site sequences with basic residue clusters (i. e. Lys cluster), near the D-site hydrophobic motif, through electrostatic interactions. To determine if p38α D-site specificity is due to its ED site, we generated ERK2 T159E/T160D (ERK2^ED^) and p38α E160T/D161T (p38α^TT^) DRS exchange mutants and examined their binding to the set of GPATCH2 and ELMSAN1 D-peptides (Fig. 5B, Fig. S6C and Table S3). Consistent with a role for the ED site in the recognition of basic D-peptides, we found that p38α^TT^ mutation led to a large decrease in binding affinity for GPATCH2-pep and ELMSAN1-pep^KxK^. Likewise, the ERK2^ED^ mutant bound GPATCH-pep about 6-fold more tightly than did WT ERK2. ERK2^ED^ mutation, however, did not promote binding to ELMSAN1-pep^KxK^, potentially because it lacks a basic residue at position 11. Furthermore, we found that while ERK2^ED^ mutation caused a large drop in affinity for ELMSAN1-pep, it had no effect on binding to GPATCH2-pep^PxP^. Finally, p38α^TT^ mutation did not promote binding to either PxP-containing peptide (ELMSAN1-pep or GPATCH-pep^PxP^). Overall, these results show that residues located in the β7-β8 turn are important for p38α recognition of basic cluster sequences, but do not contribute to ERK2 binding to the PxP (Far1) motif.

### Identification and validation of new MAPK interactors and substrates

We next examined whether D-site hits from the screens could function to recruit MAPKs in the context of their corresponding full-length proteins. We selected four proteins that were hits in both the ERK2 and p38α screens as candidate substrates: TACC1, ISG20, L3MBTL3 and ARHGEF2. All of these proteins were reportedly phosphorylated at Ser/Thr-Pro sites in phosphoproteomics studies^40^. ARHGEF2 (also called GEF-H1) is an established ERK substrate^41-43^, and L3MBTL3 was previously identified as an ERK1 binding protein in a large-scale screen^44^. We subjected purified recombinant proteins to radiolabel kinase assays in the presence of varying concentrations of a D-peptide competitor (KMT2C-D-pep) to determine whether they were phosphorylated by ERK2 or p38α in a docking-dependent manner. We found that both kinases phosphorylated TACC1 and ARHGEF2, while L3MBTL3 was phosphorylated by ERK2 alone (Fig. 6A-E), and ISG20 was not phosphorylated by either MAPK (data not shown). For each of the verified substrates, phosphorylation was reduced by the presence of the KMT2C-D-pep D-site inhibitor in a dose-dependent manner, confirming that the observed enzymatic activity is docking dependent. Taken together, these data indicate that at least some of the D-sites identified in our screens can function in substrate recruitment in an *in vitro* setting.

**Figure 6.**
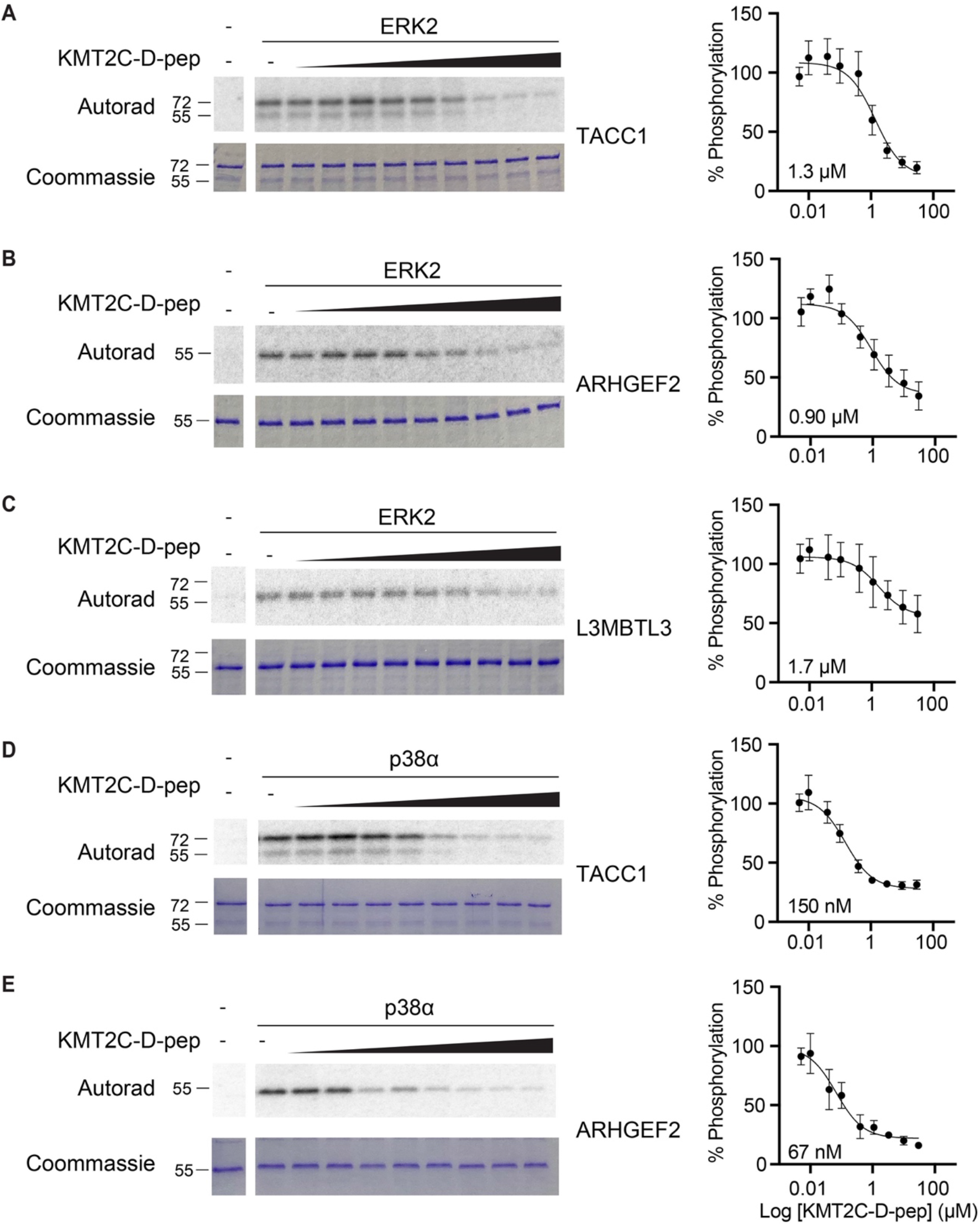
Discovery of D-site dependent ERK2 and p38α substrates. (A-E) *In vitro* radiolabel kinase assays showing phosphorylation of the indicated recombinant protein by either ERK2 or p38α. The inhibitor D-peptide (KMT2C-D-pep) was titrated in 3-fold increments from 0.5 nM to 30 µM, left to right. Quantified dose response curves for D-peptide inhibition are shown at right. Error bars indicate SD from three independent experiments.

We next turned to examine potential interactors in a cellular context. As an initial step, we started with an expanded set of seven common hit proteins to investigate whether they could interact with MAPKs in co-immunoprecipitation experiments when co-expressed in HEK293T cells with either MAPK. In these assays we found that all tested proteins co-immunoprecipitated with p38α, but that only a subset (ISG20, ARHGEF2, and MLXIPL) detectably interacted with ERK2 (Fig. 7A, B). These observations suggest that most hit D-sites promote interactions with ERK and p38 in the context of their respective full-length proteins. They further correlate with our findings from peptide competition assays that p38α tended to bind D-peptides with higher affinity than did ERK2. We next asked whether ERK or p38α could phosphorylate any of these interacting proteins in cells. HEK293T cells expressing candidate substrates were treated with EGF (with and without the MEK1/2 inhibitor trametinib) to stimulate ERK, or anisomycin (with or without the p38α/β inhibitor SB203580) to stimulate p38. As evidenced by inhibitor-sensitive mobility shift on Phos-tag SDS-PAGE, MLXIPL (also called ChREBP) was phosphorylated by both ERK and p38, and TACC1 was confirmed as a substrate of ERK2 (Fig. 7C-E). These data confirm the ability of our screening platform to discover authentic cellular MAPK substrates.

**Figure 7.**
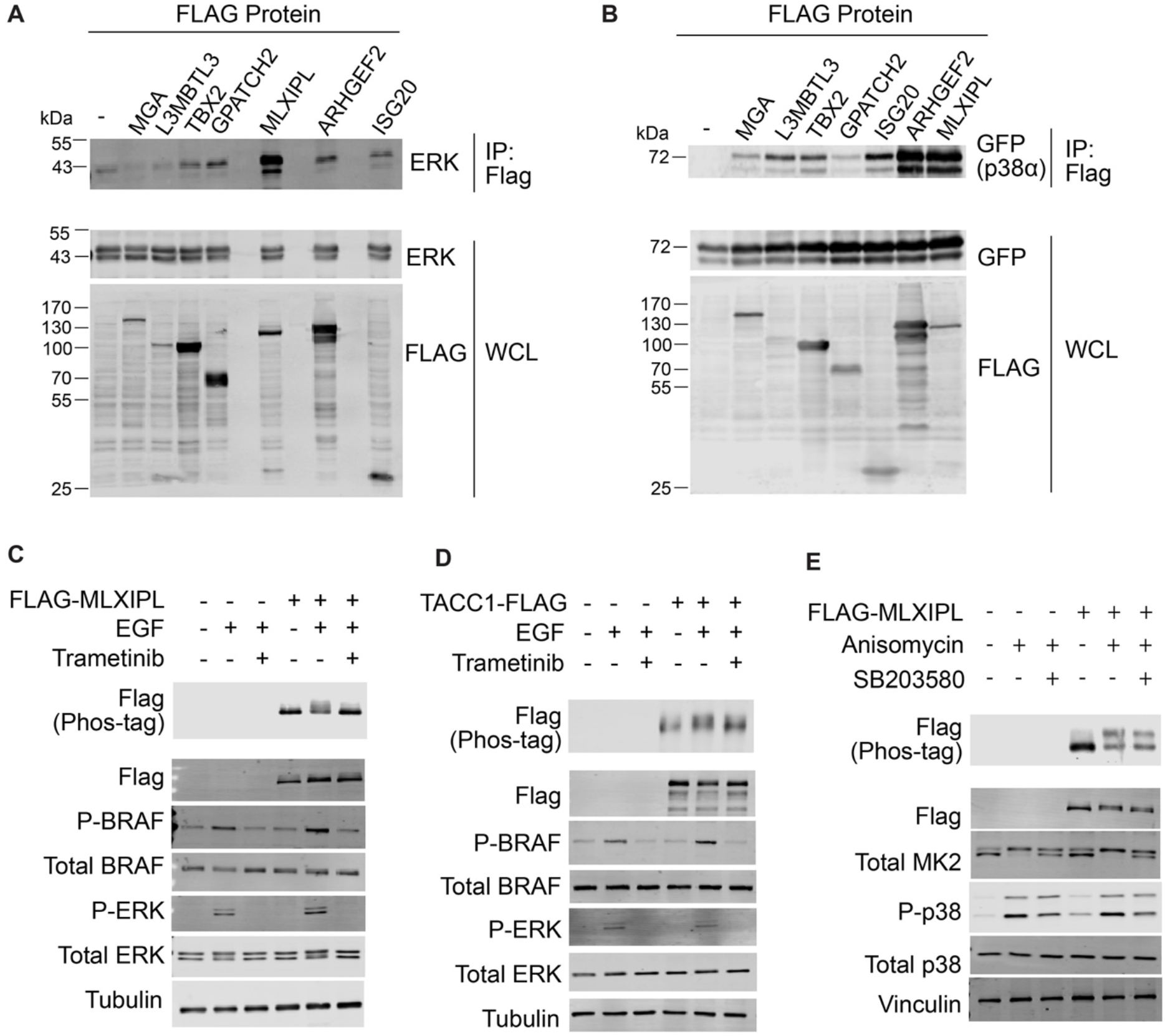
Identification of MAPK-interacting proteins and substrates in cells. (A-B) Co-immunoprecipitation of ectopically expressed ERK2 or p38α with the indicated FLAG epitope-tagged proteins. All proteins are full-length, except MGA (residues 1-1058). Experiment shown is a representative example of three independent replicates. (C-E) MAPK-dependent substrate phosphorylation in HEK293T cells. Cells were transiently transfected with the indicated FLAG epitope-tagged protein. Transfected cells were pre-treated with either 500 nM trametinib or 50 μM SB203580 as indicated for 1 hour, and then treated with 25 ng/ml EGF for 5 min to activate ERK or 500 nM anisomycin for 1 hour to activate p38. Lysates were fractionated by SDS-PAGE and immunoblotted with the indicated antibodies. Phos-tag PAGE (top slices in C-E) shows phosphorylation-dependent mobility shift of the FLAG-tagged protein. (C and E) show representatives of three independent experiments. (D) shows a representative of two independent experiments.

## DISCUSSION

Here we have reported a pooled Y2H screen to identify short proteomic sequences interacting with the MAPK DRS. Through this approach we have uncovered a large number of D-sites interacting with the MAPKs ERK2 and p38α, and we have shown that that some of these sequences function, in the context of their respective full-length proteins, as binding partners and as substrate of MAPKs in cells.

Prior work has suggested that most forward D-sites bind promiscuously to both ERK2 and p38α. Indeed, we identified the so-called MEF2A motif (P/ϕ-x-L-x-ϕ) in hits selected by both of the MAPKs^17^. However, we found that three residues in the linker region (positions 7 – 9 in our library) of the signature D-site sequence (β_2-3_-x_1-6_-ϕ-x-ϕ) can promote MAPK-selective binding. For example, ERK2 preferentially bound sequences having Pro residues located two positions upstream of the core hydrophobic motif (P-x-x-L-x-L), a feature present in several established ERK interactors including its activating kinase MAP2K1 (MEK1). In addition, we found that sequences with basic residues clustered in the linker region were highly selective for p38α. To our knowledge, this is the first example of a forward D-site motif that can strongly distinguish between these two MAPKs.

Our study shows that D-sites with higher net charge appeared to generally bind more selectively to p38α over ERK2. Interestingly, this phenomenon appeared to be driven by the presence of clustered Lys as opposed to Arg residues. Lys residues tend to increase conformational heterogeneity of IDRs ^45^ and is found more frequently than Arg both in IDRs (as annotated in the DisProt database) and in variable regions of reported linear motifs^46,47^. We speculate that the Lys cluster motif binds to p38α by forming a “fuzzy” complex, that is partly stabilized by long range electrostatic interactions with the complementary ED site and possibly the CD region of the DRS groove. By definition, a fuzzy complex occurs when one binding partner in a protein-protein complex remains dynamic, where conformational heterogeneity drive interaction specificity^48,49^. In support of this, structural studies have revealed that D-peptides adopt a variety of binding modes when interacting with MAPKs^13,15,17,23,24,27,28^, correlating well with proposed models of fuzzy complexes^50^. While forward D-sites are anchored through interactions with their C-terminal hydrophobic ϕ-x-ϕ residues, the N-terminal region can either be conformationally stable or considerably dynamic. For example, no electron density was observed for the N-terminal basic regions of the MAP2K3 and MAP2K6 D-peptides that were co-crystalized with p38α while NMR studies of these same complexes show that this region provides significant contribution to docking interaction in solution^13,15,34,37,51^. In addition, an NMR study of ERK2 bound to the substrate ETS1 suggested fuzzy interactions proximal to the D-site hydrophobic anchor^52,53^. Collectively, these studies suggest that despite making important contributions to binding affinity, clusters of basic residues in p38-selective D-sites need not adopt a single stable conformation when bound to the MAPK to mediate recruitment and specificity.

The importance of the ED site in mediating ERK versus p38 D-site specificity has been highlighted in previous studies, where mutations at this site can invert selectivity for binding to substrates harboring reverse D-sites (RSK1 and MK2/3) with more stable secondary structures^13,35^. In addition, NMR and hydrogen-deuterium exchange studies of p38/D-peptide complexes suggest that interactions involving the ED site can be more significant than those involving the CD region with peptides that have basic residues clustered N-terminal to the canonical hydrophobic motif^34,37,39,51^. This study provides evidence that the ED site can also mediate forward D-site selectivity at least in the context of ERK and p38 MAPKs. Based on the limited negative charge in the JNK DRS, we predict that this sequence feature would also confer selectivity to p38 over JNK, but additional studies will be needed to confirm if this is the case.

Our studies identified D-sites corresponding to many known MAPK binding partners and substrates^13,40,54,55^, but also identified a large number of novel sequences (Tables S1 and S2), indicating that multiple complementary approaches will be important to completely define MAPK interactomes. For example, prior research using a structure-guided *in silico* screen based on the sequence conservation and binding mode of the p38 substrate MEF2A^17^ predicted only 23% and 20% of D-sites selected respectively by ERK2 and p38α in this study (Tables S1 and S2). In addition, analysis of hits selected in our Y2H screens produced a refined definition of the MEF2A motif that includes basic residues alternating with the core hydrophobic (ϕ-x-ϕ) sequence motif. This, and other refined motifs defined in this work should facilitate more accurate computational searches to identify additional interactors that were excluded from our library. Furthermore, unbiased experimental screens such as those described here are not restricted to previously defined sequence motifs, and indeed a substantial number of our hits did not have clearly evident motifs beyond those comprising the generic D-site sequence. Our current work also identified 88% of the 71 p38α hit sequences reported in a recent study from our laboratory, in which we screened the same library for sequences inhibiting flux through the mammalian p38 pathway reconstituted in yeast^19^. Interestingly, p38 hits identified in our prior study conformed to the MEF2A class motif, while in this study we discover a substantial number of new sites that do not. While the Y2H screens identified all known MAP2K D-sites and many known substrates (Tables S1 and S2), it failed to detect some functional low affinity interactions, including D-sites from the p38α substrates MEF2A and MEF2C, that were hits in our previous screen. This suggests that D-site affinity is likely dependent on the surrounding sequence context, which differed between the two screens. Though residues surrounding the core library sequence were not predicted to be involved in direct interactions with the MAPK DRS, it has been estimated that sequence context can contribute ∼20% binding energy to SLiM-mediated interactions^56,57^. Sequence context appears particularly important when non-interacting residues reduce the entropic penalty associated with the disorder-to-order transition occurring upon binding of a SLiM to a globular domain^58^. Furthermore, a recent study reported screening a random combinatorial library of MAP2K1 D-site variants for function in promoting phosphorylation of ERK2^59^. In addition to presenting the sequence in the context of a specific ERK activator, this function-based screening approach could select for features other than simply binding to the docking groove, such as the capacity to allosterically impact ERK2 activation loop conformation or dynamics^23,37,39^.

In addition to nominating new candidate MAPK interactors, our screens have also identified previously unappreciated docking sites present in known interactions. For example, we discovered sequences corresponding to at least 16 interactors annotated in a literature-curated compendium of ERK targets (Table S1)^55^. Furthermore, we found cryptic D-sites in 18 reported p38α substrates and other interacting proteins^54^ (Table S2).

In addition, we verified that several full-length proteins harboring hit sequences interact with ERK2 or p38α in cells, and we validated the nuclear proteins MLXIPL and TACC1 as new MAPK substrates. MLXIPL encodes a glucose-activated transcription factor that upregulates lipogenic genes in the liver^60^. Phosphorylation of MLXIPL at Ser140, a Ser-Pro site located downstream of the putative docking site, had been implicated in promoting its nuclear exclusion, but kinases phosphorylating of this site were never identified^61^. Phosphorylation by MAPKs may therefore be a mechanism promoting crosstalk between signaling pathways triggered by hormones or stress with those involved in glucose sensing. TACC1 is a member of the transformation-associated coiled coil family of proteins that localize to centrosomes during mitosis, where they function to recruit chTOG/MSPS family proteins to influence microtubule dynamics^62^. Our finding that TACC1 is a docking-dependent substrate may explain observations that overexpression causes cytoplasmic sequestration of ERK2 and inhibition of downstream signaling^63^. Phosphorylation by ERK could also impact the function of TACC1 in promoting cell division and transformation^64-66^. Additional studies will be needed to determine how MAPK phosphorylation of these novel substrates impacts their function.

Overall, studies presented here expand the D-site mediated interactomes of ERK2 and p38α and provide insight into how short linear motifs mediate selectivity in MAPK signaling networks. Understanding the full repertoire of D-site dependent interactions is important in light of recent efforts to identify small molecules targeting MAPK docking grooves, which may serve as substrate-selective inhibitors^67-71^. In addition, recurrent cancer-associated mutations in the *MAPK1* gene encoding ERK2 map to the CD region and are likely to selectively disrupt phosphorylation of a subset of its substrates^72^. Our approach should also be generally applicable to provide insight into understudied MAPKs that do not participate in canonical cascades^73^, for which D-site interactions are poorly characterized. By providing a set of candidate substrates and interaction partners, this work will serve as a resource for future exploration of the regulation and function of these important kinases.

## MATERIALS AND METHODS

### Plasmids

The Y2H prey vector was generated by PCR amplification of the ELK1 transcriptional activation domain (residues 306-427) and cloning into pGAD-GH between the NcoI and PstI restriction sites, with a C-terminal V5 tag was inserted by overlap extension PCR. An AgeI restriction site was inserted downstream of Elk1 Pro326 codon to facilitate D-site sequence substitutions. Additional sequence was inserted flanking the D-site (Lys311-Pro326) to match the linker region in the oligonucleotide library. The full ORF sequence is provided in supplementary Figure S1. Elk1^ΔD^ harbors a triple Ala mutation (L319A/E320A/L321A) in the D-site motif. To construct Y2H bait vectors, rat ERK2 and p38α coding sequences were PCR-amplified and cloned into the EcoRI and SalI sites of pGBT9.

Full length cDNAs for GPATCH2, TBX2, TACC1, ISG20, L3MBTL3 and ARHGEF2 were obtained from the human ORFeome collection and shuttled into the bacterial expression plasmid pET28a by Gibson assembly or into the mammalian C-terminal 3xFLAG epitope tagged expression vector pV1900^74^ by Gateway recombination. The mammalian expression construct for mouse MLXIPL was generated by PCR-amplification from a template plasmid (from the laboratory of Isabelle Leclerc, Addgene #39235) and cloned into pcDNA3 with an N-terminal FLAG epitope tag by Gibson assembly. The mammalian expression vector for 3xFLAG-tagged human MGA produced by the laboratory of Guntram Suske was purchased from Addgene (#107715) and modified by overlap extension PCR to remove residues 1052-3065.

Bacterial expression vectors for rat ERK2 and p38α and the mammalian expression vector pCDNA3-HA-ERK2 (rat) were previously described^75^. Docking groove exchange mutants ERK2^T157E/T158D^ and p38α^E161T/D162T^ were generated by QuikChange mutagenesis. GFP-p38 produced in the laboratory of Rony Seger was purchased from Addgene (#86832). A complete list of plasmids and gene specific primers is available in supplementary Tables S4 and S5.

### Y2H Growth Assay

The Y2H host strain pJ69-4a^76^ was sequentially transformed with bait plasmids (pGBT9-GAL4BDB-ERK2 or pGBT9-GAL4DBD-p38α) and prey plasmids (pGAD-GH-GAL4AD-ELK1 variants). Overnight cultures of individual transformants in SD-Leu-Trp media were diluted to an OD_600_ of 0.1 and grown to an OD_600_ of 1.0. Five-fold serial dilutions (starting at OD_600_ = 0.5) were spotted onto agar plates with non-selective (SD-Leu-Trp) or selective (SD-Leu-Trp-His) media with 50 μM 3-amino-1,2,4-triazole [3-AT]) and incubated at 30ºC for 3-4 days.

### D-site Library Construction

The human proteomic D-site library oligonucleotide pool^19^ was amplified by 6 cycles of PCR, restriction enzyme digested and ligated into pGAD-GH-ELK1 between the SpeI and AgeI sites, replacing the native ELK1 D-site. MAX Efficiency DH10B electrocompetent *E. coli* (ThermoFisher) was transformed with the ligation reaction product. Transformed cells were spread on 15 cm Luria-Bertani Broth (LB) agar plates supplemented with 0.1 mg/ml ampicillin and grown overnight at 37°C. Plasmid DNA was isolated from pooled colonies (Qiagen Maxi prep kit). Colony counts from serial dilution plates confirmed the presence of at least 50 transformants per library sequence. The variable sequence was PCR-amplified and sequenced (Illumina HiSeq 4000) to verify clone representation of all library component. Initial clone representation was established by NGS and indicated in supplementary Dataset S1.

### Y2H Library Screen

Yeast harboring bait plasmids were transformed with the ELK1 D-site library plasmid pool and selected on agar plates containing SD-Leu-Trp media. Transformants were suspended in SD-Leu-Trp liquid media to an OD_600_ of 0.1 and grown in a shaking incubator at 30ºC. When the culture reached an OD_600_ of 1.0, it was used to inoculate 50 ml of selective media (SD-Leu-Trp-His + 50 μM 3-AT) and 50 ml of non-selective media (SD-Leu-Trp) at an OD_600_ of 0.1. Yeast cultures were subjected to two growth and dilution cycles, with 25 ml aliquots reserved for DNA extraction each time the culture reached an OD_600_ of 1.0. DNA was extracted from each aliquot and the D-site coding sequence was PCR-amplified with barcoded primers. Samples were pooled and sequenced on an Illumina HiSeq 4000 instrument with 100-bp paired end reads. Screens with ERK2 and p38α baits were performed three times independently.

Sequencing reads were mapped to internal barcodes, allowing for one base mismatch. Barcode-matched sequences were mapped to the D-site library and unique sequences were counted using the FASTX toolkit command line developed by the Hannon laboratory (https://github.com/agordon/fastx_toolkit, http://hannonlab.cshl.edu/fastx_toolkit/commandline.html). Library components with read counts below 5% of the median count in the matched control sample were discarded to avoid false positives arising from poor representation under non-selective conditions. The frequency of each D-site sequence at a given timepoint (F_Dt_) was calculated by dividing the number of reads (D) over the total read count for all sequences. An enrichment ratio for each sequence was calculated by dividing its frequency under selective conditions (F_Dt_) by its frequency under non-selective conditions (F_D0_). A plot of ln(F_Dt_/F_D0_) as a function of the number of population doublings from the linear portion of the growth curve (up to ∼15 doublings) was fit to a line, for each sequence. The enrichment score (ES) for a given sequence was defined as the slope of this line. Hits were defined as sequences with ES values greater than two standard deviations from the mean of all ES obtained under selective conditions (Z-score ≥ 2). Sequence motifs were visualized as probability logos using the pLogo algorithm^77^ and the full library as background sequences. Enrichement scores and respective Z-scores are indicated in supplementary Datasets S2 (for ERK2 screens) and S3 (for p38α screens).

### Protein Expression and Purification

His_6_-tagged ARHGEF2, TACC1 and L3MBTL3 were expressed in Rosetta DE3 pLysS *E. coli* (Invitrogen) by IPTG induction for 12-18 hours at 16ºC. Bacteria were pelleted, resuspended in lysis buffer (20 mM Tris [pH 7.5], 140 mM NaCl, 3 mM β-mercaptoethanol, 10 mM imidazole, 2 μg/μl pepstatin A, 10 μg/μl leupeptin), and lysed by sonication. Lysates were clarified by centrifugation at 3,000 x *g* and incubated with Talon® immobilized metal affinity resin (Takara #635502) for 2 hours at 4ºC. Beads were washed twice in wash buffer (20 mM Tris [pH 7.5], 150 mM NaCl, 0.01% Igepal CA-630, 10 mM imidazole), and proteins were eluted in the same buffer containing 250 mM imidazole. Proteins were dialyzed overnight in dialysis buffer (20 mM HEPES [pH 7.4], 150 mM NaCl, 10% glycerol, 1 mM DTT) and stored at -80ºC.

Active GST-tagged ERK2, ERK2^T157E/T158D^, p38α and p38α^E161T/D162T^ were produced by co-expression with His_6_-tagged hyperactive MKK constructs (pET28a-MEK1ΔN3 and pET28a-MKK6EE^75^) in BL21(DE3) *E. coli* (Invitrogen). LB cultures were grown to an OD_600_ of 0.8-1.0 before 12-hour IPTG induction at 30ºC. Cells were harvested by centrifugation and pellets were resuspended in lysis buffer (20 mM Tris [pH 7.5], 140 mM NaCl, 1 mM EDTA, 2 μg/μl pepstatin A, 10 μg/μl leupeptin, 1 mM DTT, 0.25 mg/ml lysozyme, 0.03 U/ml DNAseI, 13 mM MgCl_2_, 0.1% Igepal CA-630, 1 mM PMSF) and sonicated. Lysates, clarified as above, were incubated with glutathione Sepharose™ 4B (GE Healthcare) beads for 2 hours at 4ºC. For MAPKs to be used in peptide kinase assays, beads were washed twice in 2 ml of wash buffer (50 mM Tris [pH 8.0], 50 mM NaCl, 0.01% Igepal® CA-630, 10% glycerol, 1mM DTT) and eluted in wash buffer containing 6 mg/ml reduced glutathione. Proteins were dialyzed overnight in dialysis buffer (20 mM Tris [pH 8.0], 50 mM NaCl, 10% glycerol, 1 mM DTT) snap frozen in dry ice/ EtOH, and stored at -80ºC. For MAPKs used in radiolabel kinase assays, beads were washed once in lysis buffer and rotated at 4ºC overnight in wash buffer containing 8 U/ml thrombin (GE healthcare) to remove the GST tag. Thrombin was removed from eluate by incubation with *p*-aminobenzamidine agarose beads for 1 hour at 4ºC followed by centrifugation at 426 x *g* for 2 min. Samples were dialyzed as described above and stored at -80°C.

Recombinant proteins were analyzed by SDS-PAGE followed by Coomassie brilliant blue staining. Stained gels were scanned using Odyssey CLx imaging system (LI-COR Biosciences). Coomassie signal density was quantified using Image Studio™ Lite software (LI-COR Biosciences Version 5.2.5), and protein concentration was determined using a bovine serum albumin (BSA) standard curve.

### Immunoblotting

Cell extracts and purified proteins were fractionated by SDS-PAGE and transferred to polyvinylidene difluoride (PVDF) membrane. Membranes were blocked in 5% non-fat dry milk in TBS with 0.1% Tween (TBS-T) for 1 hour at room temperature. Primary antibodies for immunoblotting obtained from Cell Signaling Technology were against: p38 MAPK (#9212), phospho-p38 MAPK Thr180/Tyr182 (#9211), ERK1/2 (#9102), phospho-ERK1/2 Thr202/Tyr204 (E10, #9106), MK2 (#3042). Remaining primary antibodies were against: BRAF (Santa Cruz #Sc-5284), phospho-BRAF Thr753 (Invitrogen #PA5-37498), vinculin (Sigma, #V9131), α-tubulin (Sigma, #T6074), FLAG eptitope tag (M2, Sigma, #F3165), GFP (JL-8, Clontech, #632381). Secondary antibodies used included: IRDye® 800CW goat anti-mouse IgG secondary Antibody (Licor, #D10603-05) and goat anti-rabbit IgG (H+L) Highly Cross-Adsorbed Secondary Antibody, Alexa Fluor 680 (Invitrogen, #A21109). Immunoblots were imaged with Odyssey CLx imaging system (LI-COR Biosciences) and visualized with Image Studio™ Lite software (LI-COR Biosciences).

### Peptide Kinase Assays

Peptides encoding hit sequences were synthesized commercially (GenScript) with an added Tyr-Ala sequence at the N-terminus to facilitate quantification and used without further purification. Peptide stocks were prepared in DMSO and their concentrations determined from A_280_ values of aqueous dilutions calculated from the extinction coefficients of tyrosine (1280 M^-1^cm^-1^) and tryptophan (5600 M^-1^cm^-1^) residues.

Competitive peptide kinase assays were performed with a MAPK D-site-dependent reporter peptide (AssayQuant Technologies Inc, #AQT0376) following manufacturer’s instructions. Briefly, reporter peptide (5 μM) was mixed with various concentrations of unlabeled hit peptides in reaction buffer (54 mM HEPES [pH 7.4], 1.2 mM DTT, 0.01% Brij-35, 1% glycerol, 0.2 mg/ml BSA, 10 mM MgCl_2_). Kinase reactions were initiated by addition of recombinant MAPK (2 nM GST-ERK2, 4 nM GST-p38α, 20 nM GST-ERK2^T160E/T161D^ or 4 nM p38α^E161T/D162T^) and 10 μM ATP. Reactions proceeded at 30°C in a Spectra M5 plate reader equipped with SoftMax Pro Software. Florescence readings (λ_ex_ 360 nm, λ_em_ 485 nm) were collected in 90 second intervals over 21 minutes, and relative rates were calculated from the slope of the linear portion of the reaction progress curve. Data were normalized such that the rate in samples lacking competitor peptide represented 100% phosphorylation, and the lowest fluorescence value per experiment or control reactions lacking kinase represented 0% phosphorylation. Normalized slope values were plotted against peptide concentration and fitted to a sigmoidal dose-response curve using GraphPad Prism (Version 9.3.0) software to obtain IC_50_ values. Table S3 lists IC_50_ values with 95% confidence intervals for all peptides reported in this study.

### Radiolabel Kinase Assays

Radiolabel kinase assays were performed by mixing 0.5 μM His-tagged proteins with untagged MAPK (50 nM p38α or 10 nM ERK2) in reaction buffer (20 mM HEPES [pH 7.4], 10 mM MgCl_2_, 1 mM DTT, 1% DMSO) and various concentrations of KMT2C-D-pep competitor (biotin-RKRSKPKLKLKIIN-amide, synthesized at Tufts University Core Facility and HPLC purified prior to use) on ice. Kinase reactions were initiated by addition of ATP to 10 μM with 0.025 μCi/μl [γ-^32^P] ATP and immediately incubated at 30ºC for 5 min. Reactions were quenched by addition of 4x SDS-PAGE loading buffer (90 mM Tris [pH 6.8], 8% β-mercaptoethanol, 33% glycerol, 8% SDS) and heated to 95ºC for 5 min before SDS-PAGE fractionation. Gels were stained with Coomassie blue, destained, dried, and exposed to a phosphor screen. Radiolabel incorporation was visualized on a Molecular Imager FX phosphorimager (Bio-Rad) and quantified using QuantityOne Software (Bio-Rad version 11.0.5). All signals were normalized to vehicle control condition.

### Pulldown Assays

HEK293T cells were maintained in Dulbecco’s Modified Eagle Medium (DMEM) supplemented with 10% fetal bovine serum (FBS), penicillin (100 U/ml) and streptomycin (0.1 mg/ml). For co-IP experiments, cells were seeded in 10 cm plates to 60% confluence and 24 hours later were transiently co-transfected with MAPK (pcDNA3-HA-ERK2 or pEGFP-p38) and FLAG epitope-tagged substrate plasmid using polyethyleneimine (PEI). After 6 hours, cells were exchanged into fresh complete media and incubated at 37ºC for 42 hours. Cells were placed on ice, washed once with 10 ml ice cold phosphate buffered saline (PBS), suspended in 1 ml of PBS, transferred to microcentrifuge tubes and centrifuged at 106 x *g* for 5 min at 4ºC. Cell pellets were resuspended in 500 μl of lysis buffer (10 mM PIPES [pH 6.8], 50 mM NaF, 40 mM Na_4_P_2_O_7_, 50 mM NaCl, 150 mM sucrose, 0.1% Triton X-100, 2 mM Na_3_VO_4_, and cOmplete™ protease inhibitor cocktail) and kept on ice for 10 min. Lysates were cleared by centrifugation at 21,500 x *g* for 10 min and incubated with 30 μl anti-FLAG M2 affinity gel (Sigma, A2220) (pre-equilibrated in lysis buffer) for 12-16 hours at 4ºC. Beads were pelleted at 420 x *g* for 2 min at 4º, washed once with ice cold lysis buffer and twice in wash buffer (50 mM HEPES [pH 7.4], 150 mM NaCl, 5 mM β-glycerophosphate, 0.1 mM Na_3_VO_4_, 0.01% Igepal CA-630), rotating at 4ºC for 10 min per wash. Beads were resuspended in 50 μl 2X SDS-PAGE loading buffer and boiled for 5 min. Samples were filtered by centrifugation through Whatman Unifilter 0.45 μm plates (Millipore-Sigma) at 3000 x *g* for 10 minutes to remove beads. 4X SDS-PAGE loading buffer was added to filtrates and samples were boiled a second time prior to electrophoresis and immunoblotting with the indicated antibodies.

### Analysis of Protein Phosphorylation in Cells

HEK293T cells in 6-well dishes were transiently transfected with pCDNA3-FLAG-MLXIPL or pV1900-TACC1 using PEI as described above. For ERK activation, cells were starved in 0.1% FBS for 24 hours prior to addition of EGF (R&D systems #236-EG, diluted in 1 mg/ml BSA/PBS) to 25 ng/μl or carrier alone for 5 min at 37ºC. To induce p38 activation, cells were stimulated with 500 nM anisomycin or 0.001% DMSO vehicle for 1 hour at 37ºC. Where indicated cells were pre-treated with with 500 nM trametinib (SelleckChem #S2673) or 50 μM SB203580 (Sigma #559389) for 1 hour at 37ºC prior to stimulation. Following stimulation, cells were placed on ice, washed once with ice cold PBS, scraped into 200 μl of lysis buffer (20 mM Tris-HCl [pH 7.5], 150 mM NaCl, 5 mM NaF, 0.1% Triton X-100, 1 mM β-glycerophosphate, 2.5 mM Na_4_P_2_O_7_, 1 mM Na_3_VO_4_ with cOmplete™ protease inhibitor cocktail and 10 μg/ml pepstatin A), transferred to microcentrifuge tubes and rotated at 4ºC for 10 min. Lysates were centrifuged for 10 min at 21,500 x *g* and supernatant was collected. Total protein concentration was determined by Pierce BCA protein assay kit (Thermo Fisher Scientific, # 23250), samples were diluted to equal concentration, and 4X SDS-PAGE loading buffer was added. Samples were boiled at 95ºC for 5 min and fractionated on 5% SDS-polyacrylamide gels with 25 nM Phos-tag reagent (Nard Institute AAL-107) and 50 μM MnCl_2_ as reported previously^78^ or by standard SDS-PAGE, followed by immunoblotting with the indicated antibodies.

## SUPPLEMENTARY MATERIALS

Dataset S1. Elk1 D-site library variant sequences and representation

Dataset S2. Illumina sequencing data and enrichment scores for ERK2 Y2H screens

Dataset S3. Illumina sequencing data and enrichment scores for p38α Y2H screen

Table S1. ERK2 Y2H screen hits

Table S2. p38α Y2H screen hits

Table S3. D-peptide IC_50_ values from competitive inhibition kinase assays

Table S4. Complete list of plasmids

Table S5. Complete list of oligonucleotides

Figure S1. Y2H prey construct amino acid and DNA sequence

Figure S2. Competitive selection of D-site sequences in Y2H screens

Figure S3. D-peptide kinase inhibition assay dose response curves

Figure S4. Motif mining in ERK hits found by conditional probability logo analysis

Figure S5. Motif mining in p38α hits found by conditional probability logo analysis

Figure S6. Competitive inhibition dose response curves for point substituted D-peptides

## ACKNOWLEDGMENTS

We thank Titus Boggon and Lise Thomas for comments on the manuscript and members of the Turk lab for feedback on this research. We thank John Blenis and Chris Burd for providing plasmids used in this study and the Yeast Resource Center at the University of Washington for yeast two-hybrid strains.

## Author Contributions

Conceptualization: JTR, GS, BET. Methodology: JTR, BET. Investigation: JTR, BET. Visualization: JTR, BET. Supervision: BET. Writing – original draft: JTR, BET. Writing – review & editing, JTR, GS, BET.

## REFERENCES

1. Raman M, Chen W, Cobb MH. Differential regulation and properties of MAPKs. Oncogene 26, 3100–3112 (2007).

2. Lavoie H, Gagnon J, Therrien M. ERK signalling: a master regulator of cell behaviour, life and fate. Nat Rev Mol Cell Biol 21, 607–632 (2020).

3. Canovas B, Nebreda AR. Diversity and versatility of p38 kinase signalling in health and disease. Nat Rev Mol Cell Biol 22, 346–366 (2021).

4. Zeke A, Misheva M, Remenyi A, Bogoyevitch MA. JNK Signaling: Regulation and Functions Based on Complex Protein-Protein Partnerships. Microbiol Mol Biol Rev 80, 793–835 (2016).

5. Faust D, Schmitt C, Oesch F, Oesch-Bartlomowicz B, Schreck I, Weiss C, Dietrich C. Differential p38-dependent signalling in response to cellular stress and mitogenic stimulation in fibroblasts. Cell Commun Signal 10, 6 (2012).

6. Miller CJ, Turk BE. Homing in: Mechanisms of Substrate Targeting by Protein Kinases. Trends Biochem Sci 43, 380–394 (2018).

7. Johnson JL, Yaron TM, Huntsman EM, Kerelsky A, Song J, Regev A, Lin T-Y, Liberatore K, Cizin DM, Cohen BM, Vasan N, Ma Y, Krismer K, Robles JT, van de Kooij B, van Vlimmeren AE, Andrée-Busch N, Käufer N, Dorovkov MV, Ryazanov AG, Takagi Y, Kastenhuber ER, Goncalves MD, Elemento O, Taatjes DJ, Maucuer A, Yamashita A, Degterev A, Linding R, Blenis J, Hornbeck PV, Turk BE, Yaffe MB, Cantley LC. A global atlas of substrate specificities for the human serine/threonine kinome. bioRxiv, 2022.2005.2022.492882 (2022).

8. Tanoue T, Nishida E. Docking interactions in the mitogen-activated protein kinase cascades. Pharmacol Ther 93, 193–202 (2002).

9. Tanoue T, Nishida E. Molecular recognitions in the MAP kinase cascades. Cell Signal 15, 455–462 (2003).

10. Tanoue T, Adachi M, Moriguchi T, Nishida E. A conserved docking motif in MAP kinases common to substrates, activators and regulators. Nat Cell Biol 2, 110–116 (2000).

11. Peti W, Page R. Molecular basis of MAP kinase regulation. Protein Sci 22, 1698–1710 (2013).

12. Dyla M, Gonzalez Foutel NS, Otzen DE, Kjaergaard M. The optimal docking strength for reversibly tethered kinases. Proc Natl Acad Sci U S A 119, e2203098119 (2022).

13. Garai A, Zeke A, Gogl G, Toro I, Fordos F, Blankenburg H, Barkai T, Varga J, Alexa A, Emig D, Albrecht M, Remenyi A. Specificity of linear motifs that bind to a common mitogen-activated protein kinase docking groove. Sci Signal 5, ra74 (2012).

14. Bardwell AJ, Abdollahi M, Bardwell L. Docking sites on mitogen-activated protein kinase (MAPK) kinases, MAPK phosphatases and the Elk-1 transcription factor compete for MAPK binding and are crucial for enzymic activity. Biochem J 370, 1077–1085 (2003).

15. Chang CI, Xu BE, Akella R, Cobb MH, Goldsmith EJ. Crystal structures of MAP kinase p38 complexed to the docking sites on its nuclear substrate MEF2A and activator MKK3b. Mol Cell 9, 1241–1249 (2002).

16. Ho DT, Bardwell AJ, Grewal S, Iverson C, Bardwell L. Interacting JNK-docking sites in MKK7 promote binding and activation of JNK mitogen-activated protein kinases. J Biol Chem 281, 13169–13179 (2006).

17. Zeke A, Bastys T, Alexa A, Garai A, Meszaros B, Kirsch K, Dosztanyi Z, Kalinina OV, Remenyi A. Systematic discovery of linear binding motifs targeting an ancient protein interaction surface on MAP kinases. Mol Syst Biol 11, 837 (2015).

18. Whisenant TC, Ho DT, Benz RW, Rogers JS, Kaake RM, Gordon EA, Huang L, Baldi P, Bardwell L. Computational prediction and experimental verification of new MAP kinase docking sites and substrates including Gli transcription factors. PLoS Comput Biol 6, (2010).

19. Shi G, Torres Robles, J., Song, C., Salichos, L., Lou, H.J., Gerstein, M., Turk, B.E.. Proteome-wide screening for mitogen-activated protein kinase docking motifs and interactors.). Pre-print (2021).

20. Bardwell AJ, Frankson E, Bardwell L. Selectivity of docking sites in MAPK kinases. J Biol Chem 284, 13165–13173 (2009).

21. Bardwell AJ, Bardwell L. Two hydrophobic residues can determine the specificity of mitogen-activated protein kinase docking interactions. J Biol Chem 290, 26661–26674 (2015).

22. Remenyi A, Good MC, Bhattacharyya RP, Lim WA. The role of docking interactions in mediating signaling input, output, and discrimination in the yeast MAPK network. Mol Cell 20, 951–962 (2005).

23. Zhou T, Sun L, Humphreys J, Goldsmith EJ. Docking interactions induce exposure of activation loop in the MAP kinase ERK2. Structure 14, 1011–1019 (2006).

24. Liu SJ, Sun JP, Zhou B, Zhang ZY. Structural basis of docking interactions between ERK2 and MAP kinase phosphatase 3. P Natl Acad Sci USA 103, 5326–5331 (2006).

25. Akella R, Min X, Wu Q, Gardner KH, Goldsmith EJ. The third conformation of p38alpha MAP kinase observed in phosphorylated p38alpha and in solution. Structure 18, 1571–1578 (2010).

26. Ma W, Shang Y, Wei Z, Wen W, Wang W, Zhang M. Phosphorylation of DCC by ERK2 is facilitated by direct docking of the receptor P1 domain to the kinase. Structure 18, 1502–1511 (2010).

27. Zhang YY, Wu JW, Wang ZX. A distinct interaction mode revealed by the crystal structure of the kinase p38alpha with the MAPK binding domain of the phosphatase MKP5. Sci Signal 4, ra88 (2011).

28. Gogl G, Toro I, Remenyi A. Protein-peptide complex crystallization: a case study on the ERK2 mitogen-activated protein kinase. Acta Crystallogr D Biol Crystallogr 69, 486–489 (2013).

29. Kragelj J, Palencia A, Nanao MH, Maurin D, Bouvignies G, Blackledge M, Jensen MR. Structure and dynamics of the MKK7-JNK signaling complex. Proc Natl Acad Sci U S A 112, 3409–3414 (2015).

30. Zeng L, Kaoud TS, Zamora-Olivares D, Bohanon AL, Li Y, Pridgen JR, Ekpo YE, Zhuang DL, Nye JR, Telles M, Winkler M, Rivera S, Marini F, Dalby KN, Anslyn EV. Multiplexing the Quantitation of MAP Kinase Activities Using Differential Sensing. J Am Chem Soc 144, 4017–4025 (2022).

31. Yang SH, Yates PR, Whitmarsh AJ, Davis RJ, Sharrocks AD. The Elk-1 ETS-domain transcription factor contains a mitogen-activated protein kinase targeting motif. Mol Cell Biol 18, 710–720 (1998).

32. Pulido R, Zuniga A, Ullrich A. PTP-SL and STEP protein tyrosine phosphatases regulate the activation of the extracellular signal-regulated kinases ERK1 and ERK2 by association through a kinase interaction motif. Embo J 17, 7337–7350 (1998).

33. Bardwell AJ, Flatauer LJ, Matsukuma K, Thorner J, Bardwell L. A conserved docking site in MEKs mediates high-affinity binding to MAP kinases and cooperates with a scaffold protein to enhance signal transmission. J Biol Chem 276, 10374–10386 (2001).

34. Kumar GS, Page R, Peti W. The interaction of p38 with its upstream kinase MKK6. Protein Sci 30, 908–913 (2021).

35. Tanoue T, Maeda R, Adachi M, Nishida E. Identification of a docking groove on ERK and p38 MAP kinases that regulates the specificity of docking interactions. Embo J 20, 466–479 (2001).

36. Peterson LB, Yaffe MB, Imperiali B. Selective mitogen activated protein kinase activity sensors through the application of directionally programmable D domain motifs. Biochemistry 53, 5771–5778 (2014).

37. Kumar GS, Clarkson MW, Kunze MBA, Granata D, Wand AJ, Lindorff-Larsen K, Page R, Peti W. Dynamic activation and regulation of the mitogen-activated protein kinase p38. Proc Natl Acad Sci U S A 115, 4655–4660 (2018).

38. Parker BW, Goncz EJ, Krist DT, Statsyuk AV, Nesvizhskii AI, Weiss EL. Mapping low-affinity/high-specificity peptide-protein interactions using ligand-footprinting mass spectrometry. Proc Natl Acad Sci U S A 116, 21001–21011 (2019).

39. Lee T, Hoofnagle AN, Kabuyama Y, Stroud J, Min X, Goldsmith EJ, Chen L, Resing KA, Ahn NG. Docking motif interactions in MAP kinases revealed by hydrogen exchange mass spectrometry. Mol Cell 14, 43–55 (2004).

40. Hornbeck PV, Kornhauser JM, Tkachev S, Zhang B, Skrzypek E, Murray B, Latham V, Sullivan M. PhosphoSitePlus: a comprehensive resource for investigating the structure and function of experimentally determined post-translational modifications in man and mouse. Nucleic Acids Res 40, D261–270 (2012).

41. Waheed F, Dan Q, Amoozadeh Y, Zhang Y, Tanimura S, Speight P, Kapus A, Szaszi K. Central role of the exchange factor GEF-H1 in TNF-alpha-induced sequential activation of Rac, ADAM17/TACE, and RhoA in tubular epithelial cells. Mol Biol Cell 24, 1068–1082 (2013).

42. von Thun A, Preisinger C, Rath O, Schwarz JP, Ward C, Monsefi N, Rodriguez J, Garcia-Munoz A, Birtwistle M, Bienvenut W, Anderson KI, Kolch W, von Kriegsheim A. Extracellular signal-regulated kinase regulates RhoA activation and tumor cell plasticity by inhibiting guanine exchange factor H1 activity. Mol Cell Biol 33, 4526–4537 (2013).

43. Fujishiro SH, Tanimura S, Mure S, Kashimoto Y, Watanabe K, Kohno M. ERK1/2 phosphorylate GEF-H1 to enhance its guanine nucleotide exchange activity toward RhoA. Biochem Biophys Res Commun 368, 162–167 (2008).

44. Corominas R, Yang X, Lin GN, Kang S, Shen Y, Ghamsari L, Broly M, Rodriguez M, Tam S, Trigg SA, Fan C, Yi S, Tasan M, Lemmens I, Kuang X, Zhao N, Malhotra D, Michaelson JJ, Vacic V, Calderwood MA, Roth FP, Tavernier J, Horvath S, Salehi-Ashtiani K, Korkin D, Sebat J, Hill DE, Hao T, Vidal M, Iakoucheva LM. Protein interaction network of alternatively spliced isoforms from brain links genetic risk factors for autism. Nat Commun 5, 3650 (2014).

45. Zeng X, Ruff KM, Pappu RV. Competing interactions give rise to two-state behavior and switch-like transitions in charge-rich intrinsically disordered proteins. Proc Natl Acad Sci U S A 119, e2200559119 (2022).

46. Uversky VN. The intrinsic disorder alphabet. III. Dual personality of serine. Intrinsically Disord Proteins 3, e1027032 (2015).

47. Fuxreiter M, Tompa P, Simon I. Local structural disorder imparts plasticity on linear motifs. Bioinformatics 23, 950–956 (2007).

48. Sharma R, Raduly Z, Miskei M, Fuxreiter M. Fuzzy complexes: Specific binding without complete folding. Febs Lett 589, 2533–2542 (2015).

49. Tompa P, Fuxreiter M. Fuzzy complexes: polymorphism and structural disorder in protein-protein interactions. Trends Biochem Sci 33, 2–8 (2008).

50. Hadzi S, Loris R, Lah J. The sequence-ensemble relationship in fuzzy protein complexes. Proc Natl Acad Sci U S A 118, (2021).

51. Francis DM, Rozycki B, Koveal D, Hummer G, Page R, Peti W. Structural basis of p38alpha regulation by hematopoietic tyrosine phosphatase. Nat Chem Biol 7, 916–924 (2011).

52. Piserchio A, Warthaka M, Kaoud TS, Callaway K, Dalby KN, Ghose R. Local destabilization, rigid body, and fuzzy docking facilitate the phosphorylation of the transcription factor Ets-1 by the mitogen-activated protein kinase ERK2. Proc Natl Acad Sci U S A 114, E6287–E6296 (2017).

53. Piserchio A, Warthaka M, Devkota AK, Kaoud TS, Lee S, Abramczyk O, Ren P, Dalby KN, Ghose R. Solution NMR insights into docking interactions involving inactive ERK2. Biochemistry 50, 3660–3672 (2011).

54. Stark C, Breitkreutz BJ, Reguly T, Boucher L, Breitkreutz A, Tyers M. BioGRID: a general repository for interaction datasets. Nucleic Acids Res 34, D535–539 (2006).

55. Unal EB, Uhlitz F, Bluthgen N. A compendium of ERK targets. Febs Lett 591, 2607–2615 (2017).

56. Stein A, Aloy P. Contextual specificity in peptide-mediated protein interactions. PLoS One 3, e2524 (2008).

57. Ivarsson Y, Jemth P. Affinity and specificity of motif-based protein-protein interactions. Curr Opin Struct Biol 54, 26–33 (2019).

58. Flock T, Weatheritt RJ, Latysheva NS, Babu MM. Controlling entropy to tune the functions of intrinsically disordered regions. Curr Opin Struct Biol 26, 62–72 (2014).

59. Scheele RA, Lindenburg LH, Petek M, Schober M, Dalby KN, Hollfelder F. Droplet-based screening of phosphate transfer catalysis reveals how epistasis shapes MAP kinase interactions with substrates. Nat Commun 13, 844 (2022).

60. Uyeda K. Short- and Long-Term Adaptation to Altered Levels of Glucose: Fifty Years of Scientific Adventure. Annu Rev Biochem 90, 31–55 (2021).

61. Sakiyama H, Wynn RM, Lee WR, Fukasawa M, Mizuguchi H, Gardner KH, Repa JJ, Uyeda K. Regulation of nuclear import/export of carbohydrate response element-binding protein (ChREBP): interaction of an alpha-helix of ChREBP with the 14-3-3 proteins and regulation by phosphorylation. J Biol Chem 283, 24899–24908 (2008).

62. Peset I, Vernos I. The TACC proteins: TACC-ling microtubule dynamics and centrosome function. Trends Cell Biol 18, 379–388 (2008).

63. Lauffart B, Sondarva GV, Gangisetty O, Cincotta M, Still IH. Interaction of TACC proteins with the FHL family: implications for ERK signaling. J Cell Commun Signal 1, 5–15 (2007).

64. Gergely F, Karlsson C, Still I, Cowell J, Kilmartin J, Raff JW. The TACC domain identifies a family of centrosomal proteins that can interact with microtubules. P Natl Acad Sci USA 97, 14352–14357 (2000).

65. Still IH, Hamilton M, Vince P, Wolfman A, Cowell JK. Cloning of TACC1, an embryonically expressed, potentially transforming coiled coil containing gene, from the 8p11 breast cancer amplicon. Oncogene 18, 4032–4038 (1999).

66. Cully M, Shiu J, Piekorz RP, Muller WJ, Done SJ, Mak TW. Transforming acidic coiled coil 1 promotes transformation and mammary tumorigenesis. Cancer Res 65, 10363–10370 (2005).

67. Miller CJ, Muftuoglu Y, Turk BE. A high throughput assay to identify substrate-selective inhibitors of the ERK protein kinases. Biochem Pharmacol 142, 39–45 (2017).

68. Boston SR, Deshmukh R, Strome S, Priyakumar UD, MacKerell AD, Jr., Shapiro P. Characterization of ERK docking domain inhibitors that induce apoptosis by targeting Rsk-1 and caspase-9. BMC Cancer 11, 7 (2011).

69. Sammons RM, Perry NA, Li Y, Cho EJ, Piserchio A, Zamora-Olivares DP, Ghose R, Kaoud TS, Debevec G, Bartholomeusz C, Gurevich VV, Iverson TM, Giulianotti M, Houghten RA, Dalby KN. A Novel Class of Common Docking Domain Inhibitors That Prevent ERK2 Activation and Substrate Phosphorylation. ACS Chem Biol 14, 1183–1194 (2019).

70. Alexa A, Ember O, Szabo I, Mo’ath Y, Poti AL, Remenyi A, Banoczi Z. Peptide Based Inhibitors of Protein Binding to the Mitogen-Activated Protein Kinase Docking Groove. Front Mol Biosci 8, 690429 (2021).

71. Samadani R, Zhang J, Brophy A, Oashi T, Priyakumar UD, Raman EP, St John FJ, Jung KY, Fletcher S, Pozharski E, MacKerell AD, Jr., Shapiro P. Small-molecule inhibitors of ERK-mediated immediate early gene expression and proliferation of melanoma cells expressing mutated BRaf. Biochem J 467, 425–438 (2015).

72. Taylor CAt, Cormier KW, Keenan SE, Earnest S, Stippec S, Wichaidit C, Juang YC, Wang J, Shvartsman SY, Goldsmith EJ, Cobb MH. Functional divergence caused by mutations in an energetic hotspot in ERK2. Proc Natl Acad Sci U S A 116, 15514–15523 (2019).

73. Coulombe P, Meloche S. Atypical mitogen-activated protein kinases: structure, regulation and functions. Biochim Biophys Acta 1773, 1376–1387 (2007).

74. Miller CJ, Lou HJ, Simpson C, van de Kooij B, Ha BH, Fisher OS, Pirman NL, Boggon TJ, Rinehart J, Yaffe MB, Linding R, Turk BE. Comprehensive profiling of the STE20 kinase family defines features essential for selective substrate targeting and signaling output. PLoS Biol 17, e2006540 (2019).

75. Sheridan DL, Kong Y, Parker SA, Dalby KN, Turk BE. Substrate discrimination among mitogen-activated protein kinases through distinct docking sequence motifs. J Biol Chem 283, 19511–19520 (2008).

76. James P, Halladay J, Craig EA. Genomic libraries and a host strain designed for highly efficient two-hybrid selection in yeast. Genetics 144, 1425–1436 (1996).

77. O’Shea JP, Chou MF, Quader SA, Ryan JK, Church GM, Schwartz D. pLogo: a probabilistic approach to visualizing sequence motifs. Nat Methods 10, 1211–1212 (2013).

78. Kinoshita E, Kinoshita-Kikuta E, Koike T. Separation and detection of large phosphoproteins using Phos-tag SDS-PAGE. Nat Protoc 4, 1513–1521 (2009).

79. Goddard TD, Huang CC, Meng EC, Pettersen EF, Couch GS, Morris JH, Ferrin TE. UCSF ChimeraX: Meeting modern challenges in visualization and analysis. Protein Sci 27, 14–25 (2018).

