## Supplemental Figures for "Linear motif specificity in signaling through p38α and ERK2 mitogen-activated protein kinases"

| | | | | | | | | | | | | | | | | |
|---|---|---|---|---|---|---|---|---|---|---|---|---|---|---|---|---|
| M | D | K | A | E | L | I | P | E | P | P | K | K | K | R | K | V |
| ATG | GAT | AAA | GCG | GAA | TTA | ATT | CCC | GAG | CCT | CCA | AAA | AAG | AAG | AGA | AAG | GTC |
| E | L | G | T | A | A | N | F | N | Q | S | G | N | I | A | D | S |
| GAA | TTG | GGT | ACC | GCC | GCC | AAT | TTT | AAT | CAA | AGT | GGG | AAT | ATT | GCT | GAT | AGC |
| S | L | S | F | T | F | T | N | S | S | N | G | P | N | L | I | T |
| TCA | TTG | TCC | TTC | ACT | TTC | ACT | AAC | AGT | AGC | AAC | GGT | CCG | AAC | CTC | ATA | ACA |
| T | Q | T | N | S | Q | A | L | S | Q | P | I | A | S | S | N | V |
| ACT | CAA | ACA | AAT | TCT | CAA | GCG | CTT | TCA | CAA | CCA | ATT | GCC | TCC | TCT | AAC | GTT |
| H | D | N | F | M | N | N | E | I | T | A | S | K | I | D | D | G |
| CAT | GAT | AAC | TTC | ATG | AAT | AAT | GAA | ATC | ACG | GCT | AGT | AAA | ATT | GAT | GAT | GGT |
| N | N | S | K | P | L | S | P | G | W | T | D | Q | T | A | Y | N |
| AAT | AAT | TCA | AAA | CCA | CTG | TCA | CCT | GGT | TGG | ACG | GAC | CAA | ACT | GCG | TAT | AAC |
| A | F | G | I | T | T | G | M | F | N | T | T | T | M | D | D | V |
| GCG | TTT | GGA | ATC | ACT | ACA | GGG | ATG | TTT | AAT | ACC | ACT | ACA | ATG | GAT | GAT | GTA |
| Y | N | Y | L | F | D | D | E | D | T | P | P | N | P | K | K | E |
| TAT | AAC | TAT | CTA | TTC | GAT | GAT | GAA | GAT | ACC | CCA | CCA | AAC | CCA | AAA | AAA | GAG |
| | | | SpeI | | | | | | | | | | | | | |
| I | L | E | L | V | G | R | G | S | M | S | Q | x | x | x | x | x |
| ATC | CTA | GAA | CTA | GTG | GGT | CGC | GGA | TCT | ATG | TCT | CAG | NNN | NNN | NNN | NNN | NNN |
| x | x | x | x | x | x | x | x | x | E | A | F | E | Q | P | Q | H |
| NNN | NNN | NNN | NNN | NNN | NNN | NNN | NNN | NNN | GAA | GCT | TTT | GAA | CAA | CCT | CAG | CAC |
| AgeI | | | | | | | | | | | | | | | | |
| T | G | S | L | L | G | G | Q | G | P | E | R | T | P | G | S | G |
| ACC | GGT | AGC | CTG | CTG | GGT | GGC | CAG | GGA | CCT | GAA | CGG | ACT | CCA | GGA | TCA | GGA |
| T | S | S | G | L | Q | A | P | G | P | A | L | T | P | S | L | L |
| ACA | AGC | TCT | GGT | CTT | CAG | GCA | CCG | GGG | CCA | GCG | CTA | ACG | CCA | TCC | CTG | CTC |
| P | T | H | T | L | T | P | V | L | L | T | P | S | S | L | P | P |
| CCC | ACA | CAT | ACC | TTG | ACC | CCG | GTG | CTG | CTG | ACA | CCC | AGC | TCG | CTG | CCC | CCT |
| S | I | H | F | W | S | T | L | S | P | I | A | P | R | S | P | A |
| AGC | ATC | CAT | TTC | TGG | AGC | ACT | CTG | AGT | CCA | ATT | GCA | CCC | CGT | AGT | CCA | GCC |
| K | L | S | F | Q | F | P | S | S | G | S | A | Q | V | H | I | P |
| AAG | CTC | TCC | TTC | CAG | TTT | CCG | TCC | AGT | GGC | AGC | GCA | CAG | GTG | CAC | ATC | CCT |
| S | I | V | D | G | L | S | T | P | V | V | L | S | P | G | P | Q |
| TCC | ATC | GTG | GAT | GGC | CTC | TCG | ACC | CCC | GTG | GTG | CTC | TCC | CCA | GGG | CCC | CAG |
| G | G | G | A | A | A | G | K | P | I | P | N | P | L | L | G | L |
| GGT | GGA | GGT | GCT | GCA | GCT | GGA | AAG | CCT | ATC | CCT | AAC | CCT | CTC | CTC | GGT | CTC |
| D | S | T | * | | | | | | | | | | | | | |
| GAT | TCT | ACG | TAA | | | | | | | | | | | | | |

**Figure S1. Y2H prey construct amino acid and DNA sequence**. DNA sequence and translation of prey fusion encoded in the pGAD GH-ELK1 plasmid used D-site library cloning and Y2H screens. Gal4 activation domain (italics), ELK1 transcriptional activation domain (bold) with the variable D-site region (purple), restriction sites (blue), flanking sites for PCR amplification (green) and V5 tag (Grey) are shown.

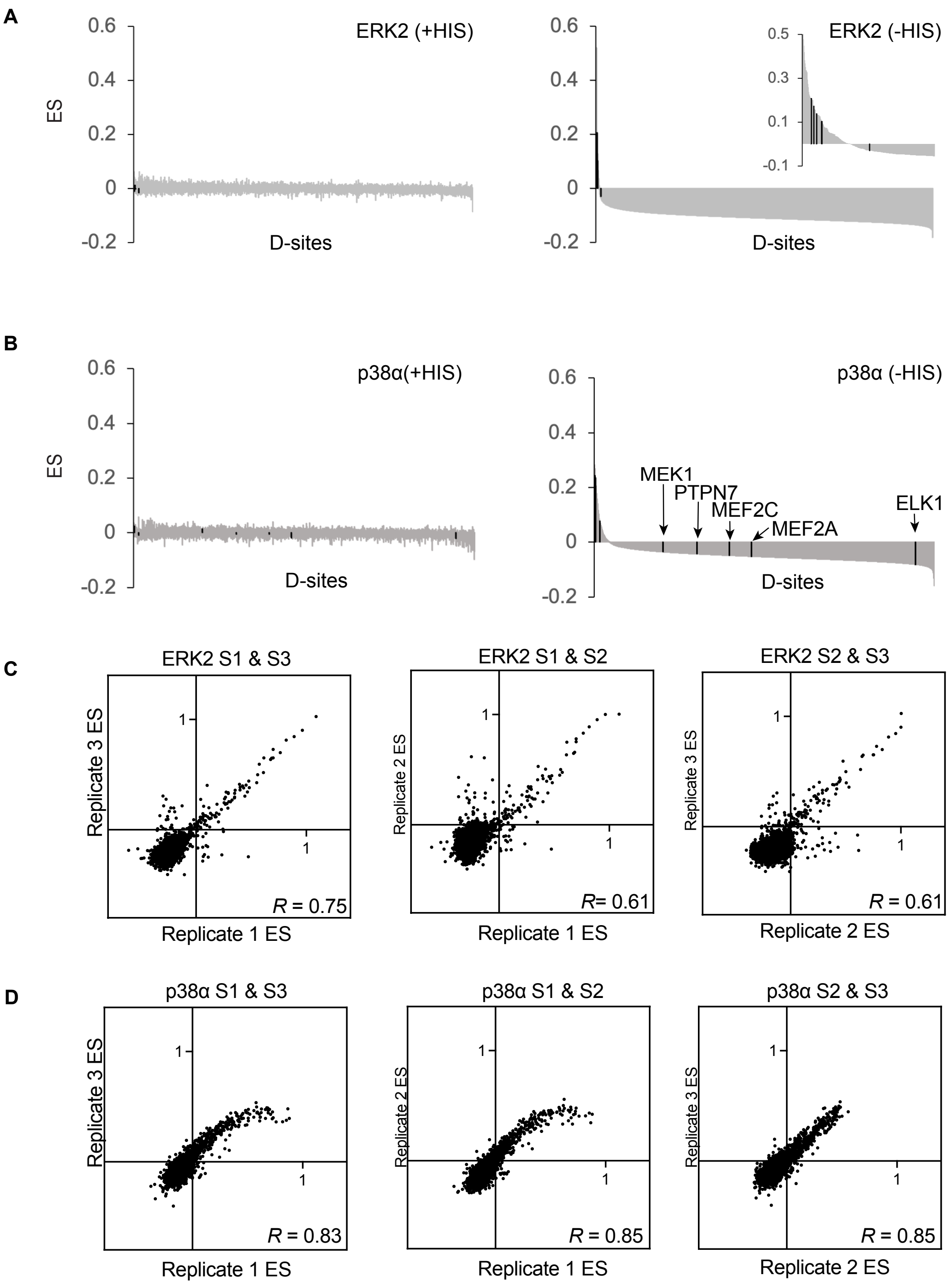


**Figure S2. Competitive selection of D-site sequences in Y2H Screens.** Waterfall plots showing enrichment scores (ES) for each D-site in the (A) ERK2 or (B) p38$\alpha$ screens under non-selective (+HIS, left panel) and selective conditions (-HIS, right panel). D-sites are sorted in descending order by ES under selective conditions for the respective MAPK. (C-D) Individual ES correlations (Pearson *R* shown) for each pairwise combination of three independent replicate screens (S) with (C) ERK2 and (D) p38$\alpha$ under selective conditions.

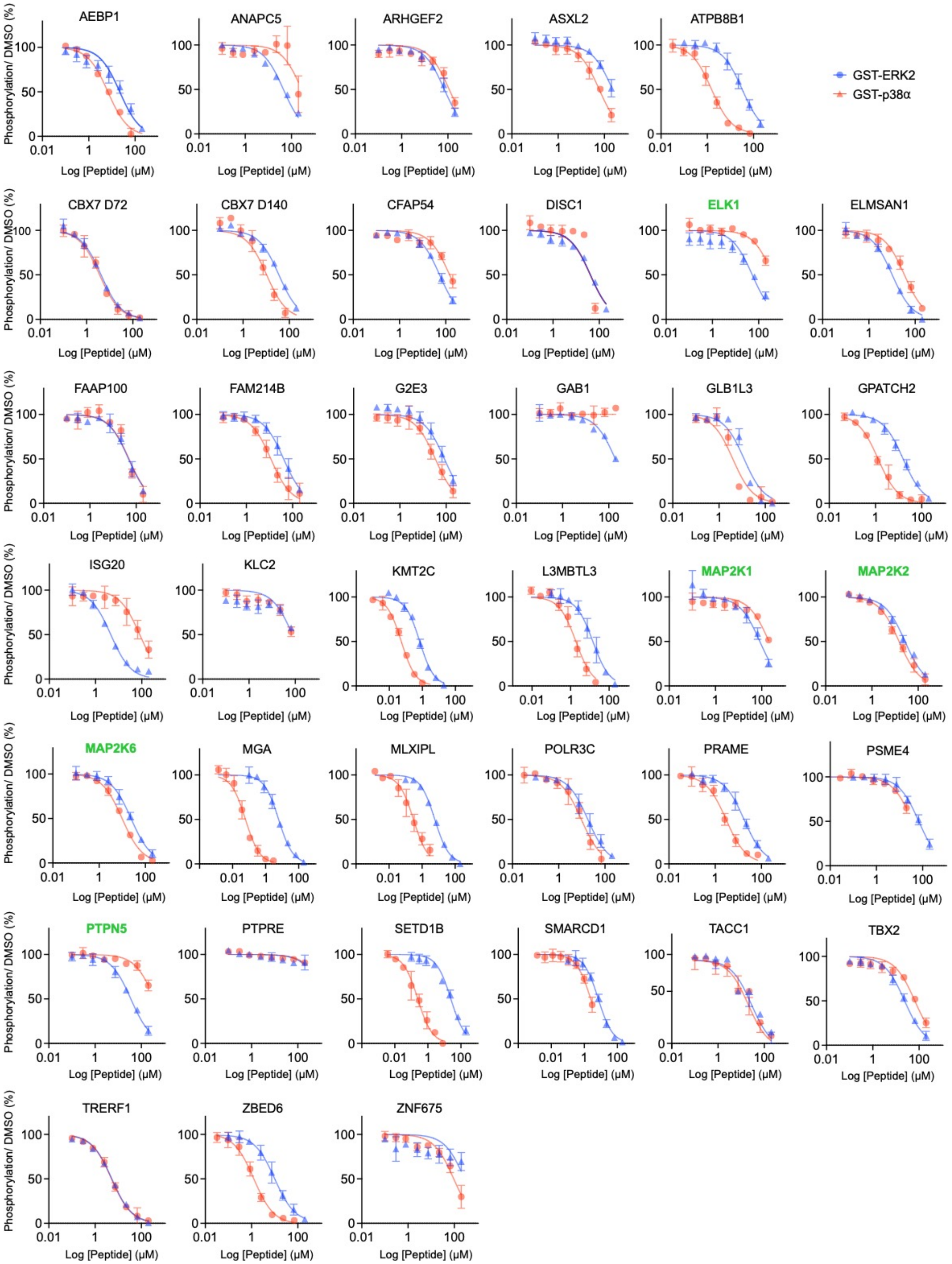


**Figure S3. D-peptide kinase inhibition assay dose response curves.** Data points represent the mean from three replicate experiments. Error bars represent SD. Previously known D-sites are shown in green for reference.

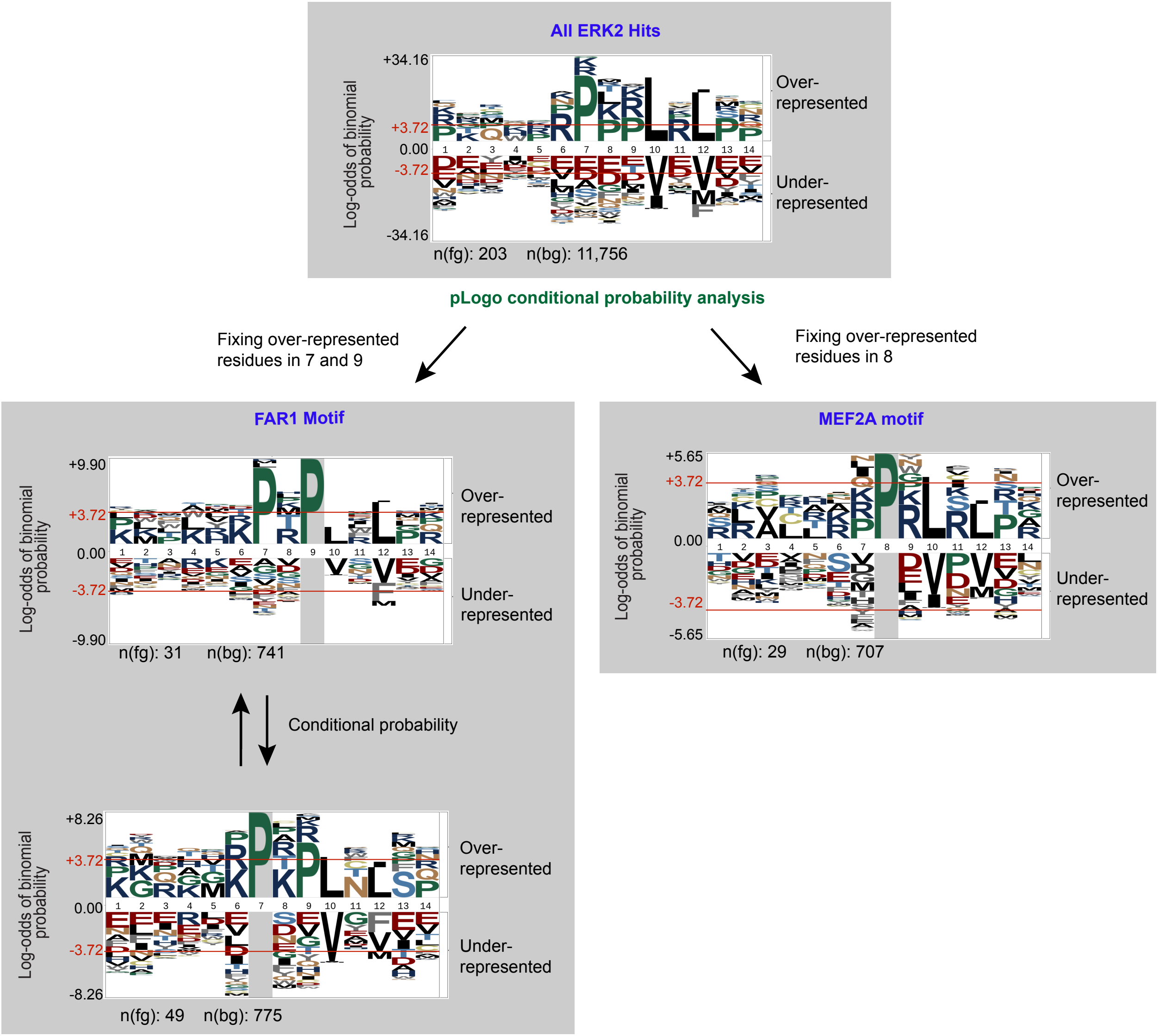


**Figure S4. Motif mining in ERK2 hits found by conditional probability analysis**. Top panel, pLogo derived from multiple sequence alignment of all 203 sequences with Z-scores ≥ 2 using all library sequences as the background. Unbiased identification of the the FAR1-type by independently fixing Pro at position 7 or 9 (left panel) and MEF2A class motif by fixing Pro at position 8 (right panel).

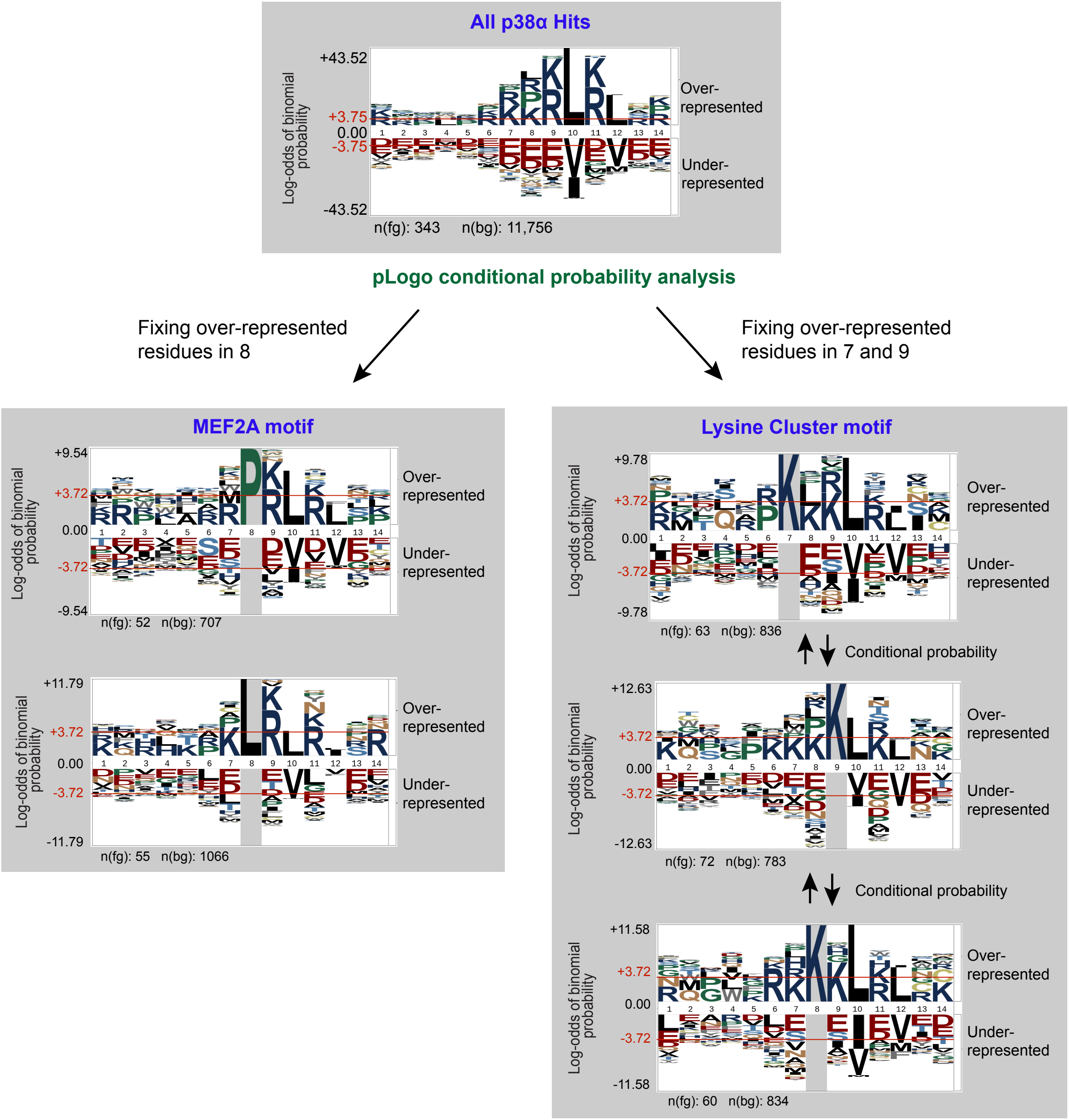


**Figure S5. Motif mining in p38α hits found by conditional probability analysis.** Top panel, pLogo analysis of the all 343 p38α hits on the background of the full library sequence. Left panels, fixing Leu or Pro residues in position 8 returns a modified version of the known MEF2A motif with particular over-representation for basic residues in position 9 and 11. Right panel, Lys cluster motif is identified by conditionally over-represented Lys residues, significantly selected at positions 7-9.

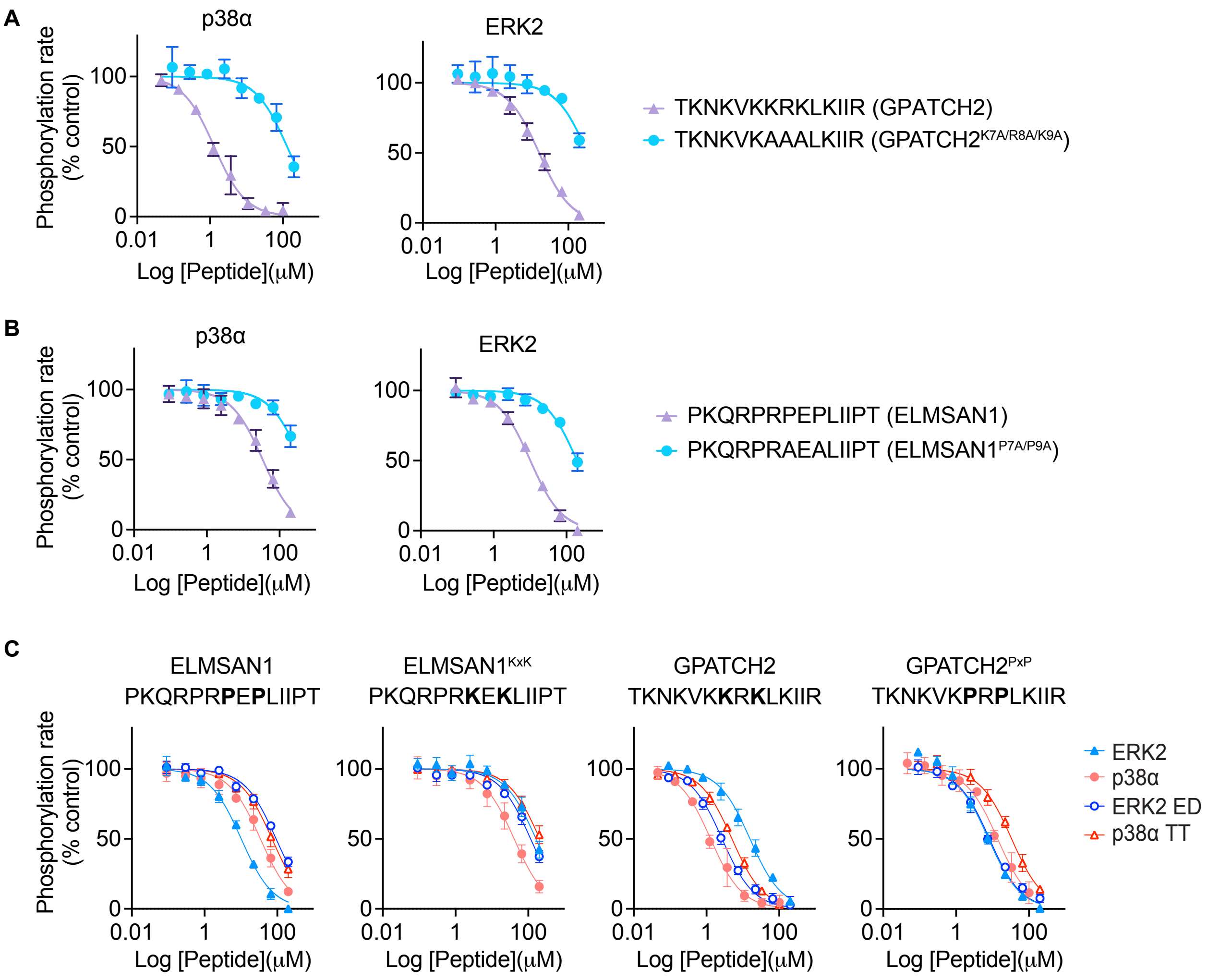


**Figure S6. Competitive inhibition dose response curves for point substituted D-peptides.** (A-B) Dose response curves for inhibition of ERK2 (blue, triangle) or p38α (orange, circles) by (A) GPATCH2$^{WT}$ or GPATCH2$^{K7A/R8A/K9A}$ and (B) ELMSAN1$^{WT}$ or ELMSAN1$^{P7A/P9A}$. (C) Dose response curves for inhibition of ERK2 (blue, triangle), p38α (orange, circle) or their respective ED region exchange mutants ERK2$^{T159E/T160D}$ (ERK2 ED; blue, open circle) or p38α$^{E160T/D161T}$ (p38α TT; red, open trianlge) by the indicated synthetic D-peptides. Data points show the mean from three independent experiments. Error bars represent SD.
